# iAPEX: Improved APEX-based proximity labeling for subcellular proteomics using an enzymatic reaction cascade

**DOI:** 10.1101/2025.01.10.632381

**Authors:** Tommy J. Sroka, Lea K. Sanwald, Avishek Prasai, Josefine Hoeren, Karina von der Malsburg, Valerie Chaumet, Per Haberkant, Kerstin Feistel, David U. Mick

## Abstract

Ascorbate peroxidase (APEX) is a versatile labeling enzyme used for live-cell proteomics at high spatial and temporal resolution. However, toxicity of its substrate hydrogen peroxide and background labeling by endogenous peroxidases limit its use to *in vitro* studies of specific cell types. By combining APEX2 with a D-amino acid oxidase to locally produce hydrogen peroxide, we establish a more versatile, improved APEX (iAPEX) workflow that minimizes hydrogen peroxide toxicity and reduces non-specific background labeling. We employ iAPEX to perform live-cell proteomics of a cellular microdomain, the primary cilium, in previously inaccessible cell lines, leading to the identification of new ciliary proteins. Our study robustly validates common ciliary proteins across two distinct cell lines, while observed differences may reflect heterogeneity in primary cilia proteomes. iAPEX proximity labeling in *Xenopus laevis* provides a proof-of-concept for future *in vivo* applications.

## INTRODUCTION

Proximity labeling technologies provide the biological and chemical sciences new applications to study nucleic acids, lipids and proteins^1–4^. Conceptually, proximity labeling uses enzymes fused to proteins of interest or directed to specific locations to determine their molecular environments by purifying proximity labeled biomolecules. Combining this approach with mass spectrometry-based proteomics allows unbiased and systematic determination of transient protein interactions as well as (sub)proteomes of organelles or cellular microcompartments that are inaccessible to biochemical purification methods^5,6^.

The primary cilium is a solitary plasma membrane microdomain with important functions in developmental biology and tissue maintenance^7,8^. It functions as a specialized signaling compartment that translates extracellular cues into cellular responses by intricate mechanisms, employing second messengers and dynamic protein transport to and from the primary cilium^9,10^. Defects in these processes have been implicated in syndromic disorders, termed ciliopathies, affecting several tissues and cell types^7^. A current hypothesis posits that cell type-dependent differences in the composition of primary cilia account for the pleiotropic syndromic disorders, as cilia dysfunctions in specific signal transduction mechanisms may have cell type- and tissue-specific consequences. Due to its small size (∼1:10,000^th^ of the cell)^11^ and difficulty to isolate pure primary cilia by classic biochemical methods^12^, proximity labeling approaches have been utilized to determine primary cilia proteomes of disease models and to investigate basic cilia biology^13–16^, such as dissecting the molecular composition of primary cilia during active signaling^17,18^. However, our knowledge of the protein composition of primary cilia is still incomplete as it stems from very few model cell types that are amenable to the available technologies.

While new proximity labeling technologies are an active area of research^19^, the most frequently used proximity labeling methods are based on two enzymatic activities that use different chemistries to label nearby proteins: 1) promiscuous biotin ligases, such as BioID, or 2) peroxidases, such as ascorbate peroxidase (APEX). BioID-based technologies are simple to use and only require biotin and ATP as substrates. In cases where the biological system requires a constant supply of biotin, BioID is continuously active and consumes biotin, resulting in persistent labeling -a major challenge for time-resolved studies and *in vivo* application^20^. A recently developed light-activatable variant, LOV-turbo^21^, is a remarkable improvement, however, comes at the cost of a more complex experimental setup. APEX-based approaches require two substrates, a tyramide is oxidized to produce a phenoxyl radical that reacts with nearby targets, while hydrogen peroxide (H_2_O_2_) is reduced to water^22^. H_2_O_2_ supplementation provides control over the enzymatic activity, yet, endogenous cellular peroxidases can also use H_2_O_2_ to oxidize tyramides^2,23^. To account for such potential non-specific labeling, experimental designs include complex, time- and resource-consuming specificity controls, such as mislocalized APEX transgenes or genetic ablation of the target structure^24^. Most critically, APEX requires H_2_O_2_ to be supplied in high concentrations (mM), which induces oxidative damage in virtually all biological contexts posing a significant challenge for *in vivo* studies^25–27^.

Here, we show that many commonly used cell culture models are incompatible with previous APEX2-based proximity labeling methods. Undesired background often exceeds APEX2-mediated proximity biotinylation due to endogenous peroxidase activities when potentially toxic H_2_O_2_ is added externally. In the work presented here, we could overcome these limitations of APEX-based proximity labeling by employing the enzyme D-amino acid oxidase (DAAO) from *Rhodotorula gracilis*^28^ to locally generate H_2_O_2_. Thereby, APEX2-mediated biotinylation is rendered dependent on an enzyme cascade, yielding a more versatile and improved APEX (iAPEX) system, which 1) expands the applicability to additional biological systems, 2) reduces toxicity by avoiding addition of exogenous H_2_O_2_, and 3) increases specificity of APEX labeling to circumvent complex genetic controls. Using this methodology, we could successfully determine the proteomes of primary cilia of cell types hitherto inaccessible to conventional APEX proximity labeling and provide proof-of-concept for the *in vivo* application of iAPEX in *Xenopus leavis*.

## RESULTS

### D-amino acid oxidase can activate ascorbate peroxidase

Quantitative proteomics on subcellular microdomains is technically challenging. Since proteomic information on primary cilia is limited to few cell types, we aimed to determine cilia proteomes of cell lines commonly used to study primary cilia by employing cilia-APEX2, an experimental setup we have successfully applied to study the ciliary proteome in a quantitative and time-resolved manner in kidney epithelial cells^17^. As APEX2-based proximity labeling is widespread, we envisioned an easy transfer of the methodology to other cell types. Yet, performing APEX2 labeling reactions using hydrogen peroxide (H_2_O_2_) resulted in various degrees of background biotinylation within cell types of interest, such as C2C12 myoblasts, 3T3-L1 pre-adipocytes, and NIH/3T3 fibroblasts (**Fig. 1A**). Even in the absence of an APEX2 enzyme, biotinylation throughout the cell body of these cells exceeded the levels observed in primary cilia in cilia-APEX2 expressing IMCD3 cells. After generating an NIH/3T3 cell line that stably expresses cilia-APEX2, we performed APEX2 labeling reactions by H_2_O_2_ addition and investigated the amounts of biotinylation by SDS-PAGE and Western Blotting (**Fig. 1B**). Surprisingly, biotinylation in NIH/3T3 cells was independent of the presence of the cilia-APEX2 enzyme and greatly surpassed the amounts observed in the well-established IMCD3 cell line (**Fig. 1B**, lanes 4 *vs.* 6), which indicated excessive background biotinylation by endogenous peroxidases.

**Fig. 1:**
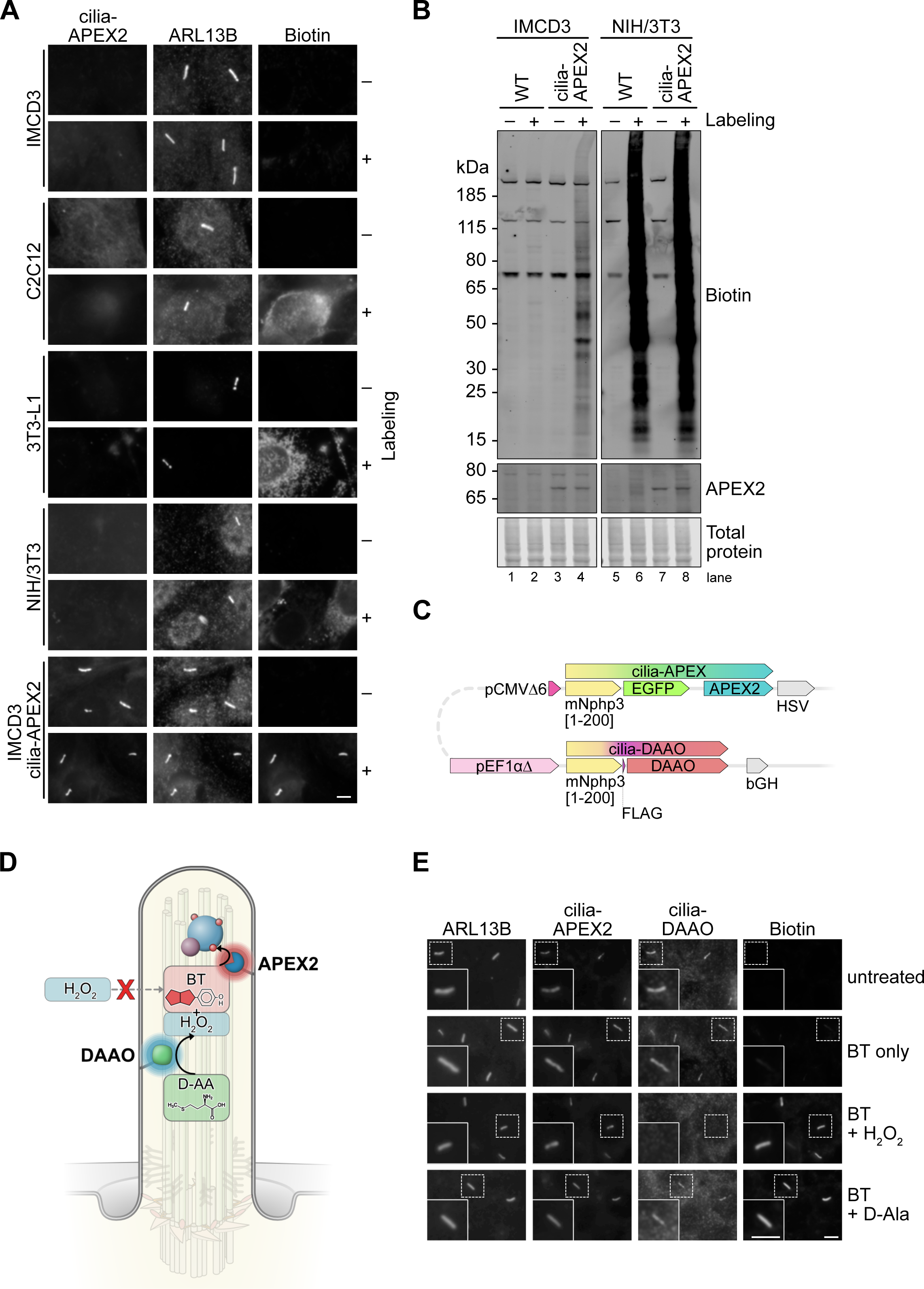
Background biotinylation in various cell types limits APEX-based proximity labeling applications. **(A)** Immunofluorescence micrographs of IMCD3, C2C12, 3T3-L1, and NIH/3T3 cells, including IMCD3 cells stably expressing cilia-APEX2. Cells were left untreated (–) or subjected to APEX2 proximity labeling (+). For the latter cells were incubated with 500 µM biotin tyramide (BT) for 30 min followed by 1 mM hydrogen peroxide (H_2_O_2_) for 3 min. Post fixation, primary cilia were visualized by ARL13B antibody staining. Biotin was detected by fluorescently labeled streptavidin. **(B)** Wild-type (WT) and cilia-APEX2-expressing IMCD3 and NIH/3T3 cells were lysed before (–) or after APEX2 proximity labeling (+), and 20 µg of protein analyzed by SDS-PAGE and Western Blotting. Biotinylated proteins were visualized by fluorescent streptavidin, equal protein loading confirmed by total protein stain. **(C)** Diagram of cilia-iAPEX expression cassette with cilia-APEX2 and cilia-DAAO transgenes in a head-to-head orientation. Cassette is part of a dual-expression vector designed for targeted Flp-In recombinase-mediated stable genomic integration into cell lines containing a Flp recombination target (FRT) site. The vector allows a low-level expression of cilia-APEX2 and cilia-DAAO from a truncated cytomegalovirus promoter (pCMVΔ6) and a EF1α promoter lacking the TATA box (pEF1αΔ), respectively . Both enzymes are fused to an N-terminal cilia-targeting sequence (the first 200 amino acids of murine Nephrocystin-3 (mNphp3)) and carry detection tags—enhanced GFP (eGFP) for APEX2 and FLAG for DAAO. **(D)** Schematic of a primary cilium harboring the improved APEX (iAPEX) proximity labeling technology. APEX2 and D-amino acid oxidase (DAAO) are genetically targeted to primary cilia using constructs displayed in (C). A D-amino acid (D-AA) serves as a DAAO substrate for *in situ* hydrogen peroxide (H_2_O_2_) production through oxidative deamination. Locally produced H_2_O_2_ activates nearby APEX2 to use biotin tyramide (BT) as a substrate for proximity biotinylation of nearby proteins, which overcomes the need for external H_2_O_2_ addition (red X). **(E)** Representative immunofluorescence micrographs showing APEX2 proximity labeling in primary cilia of IMCD3 cells stably expressing the cilia-targeted iAPEX enzyme cascade. Cilia-APEX2 is detected by GFP autofluorescence, cilia-DAAO by anti-FLAG antibodies. For classic APEX2 proximity labeling cells were incubated with biotin tyramide (BT, 500 µM) for 30 min and H_2_O_2_ for 3 min. For DAAO-facilitated proximity labeling, D-alanine (D-Ala, 10 mM) was added during BT incubation for 30 min.

To overcome non-specific proximity labeling and avoid external addition of H_2_O_2_ we expressed D-amino acid oxidase (DAAO) from *Rhodotorula gracilis* that oxidizes D-amino acids, the rare enantiomers of the predominant L-amino acids, to produce H_2_O_2_ intracellularly^28–31^. To specify and restrict the subcellular localization of H_2_O_2_ production we targeted DAAO to primary cilia by fusing it to the first 200 amino acids of the ciliary protein NPHP3, which we term cilia-DAAO (**Fig. 1C**). We hypothesized that locally produced H_2_O_2_ would be immediately consumed by nearby APEX2 to oxidize biotin tyramide for proximity labeling (**Fig. 1D**). To confirm the functionality of this enzymatic cascade in primary cilia, we generated an IMCD3 cell line stably expressing both cilia-APEX2 and cilia-DAAO, which localize specifically to primary cilia (**Fig. 1E**). In this cell line, proximity biotinylation in primary cilia could be achieved in the presence of biotin tyramide either by addition of H_2_O_2_ (to activate APEX2 directly) or by providing the DAAO substrate D-alanine (D-Ala) (**Fig. 1E**). Although small molecules can diffuse freely between the cilium and the cytoplasm^32^, the functionality of the cascade required DAAO to be localized to cilia, as a DAAO enzyme localized to the cytosol (cyto-DAAO) did not activate cilia-APEX2 (**Fig. S1A**). This suggests that H_2_O_2_ produced by DAAO in the cytoplasm does not diffuse into primary cilia, probably due to rapid detoxification. Interestingly, in cilia with strong biotin signals we observed a reduction in the cilia-DAAO signal (**Fig. S1B-C**). As cilia-DAAO is detected *via* the FLAG epitope, we interpret this anti-correlation as a potential biotinylation of the tyrosine residue within the FLAG epitope, which may mask antibody binding.

D-amino acids are inert in most biological systems but show biological activity in rare instances, such as D-serine as a putative gliotransmitter^33–35^. Therefore, we tested different amino acids and derivatives as potential DAAO substrates for iAPEX-based proximity labeling in the cilia-APEX2 and cilia-DAAO expressing IMCD3 cell line. Except for D-serine and D-valine, all D-amino acids tested induced biotin tyramide-dependent biotinylation in primary cilia in IMCD3 cells, which confirmed suitability and stereo-specificity of these substrates for iAPEX labeling (**Fig. S1D**). Taken together, our experiments confirmed the functionality of the DAAO-APEX enzymatic cascade, which we term “improved APEX” (“iAPEX”) proximity labeling.

### Local hydrogen peroxide production minimizes toxicity

As the local restriction of APEX2 activation within primary cilia may limit its application for whole cilium proteomics we assessed the subciliary localization of biotinylated proteins by ultrastructure expansion microscopy (U-ExM)^36^. U-ExM confirmed that both cilia-APEX2 and cilia-DAAO were confined to the membrane of the primary cilium (**Fig. 2A**) with varying degrees of co-localization. However, after activation of the iAPEX cascade, biotinylation was not restricted to the membrane and could be detected throughout the entire cilium, indistinguishable from the activation by external addition of H_2_O_2_ (**Fig. 2B**), indicating that iAPEX labeling generates sufficient phenoxyl radicals to probe the entire cilium.

**Fig. 2:**
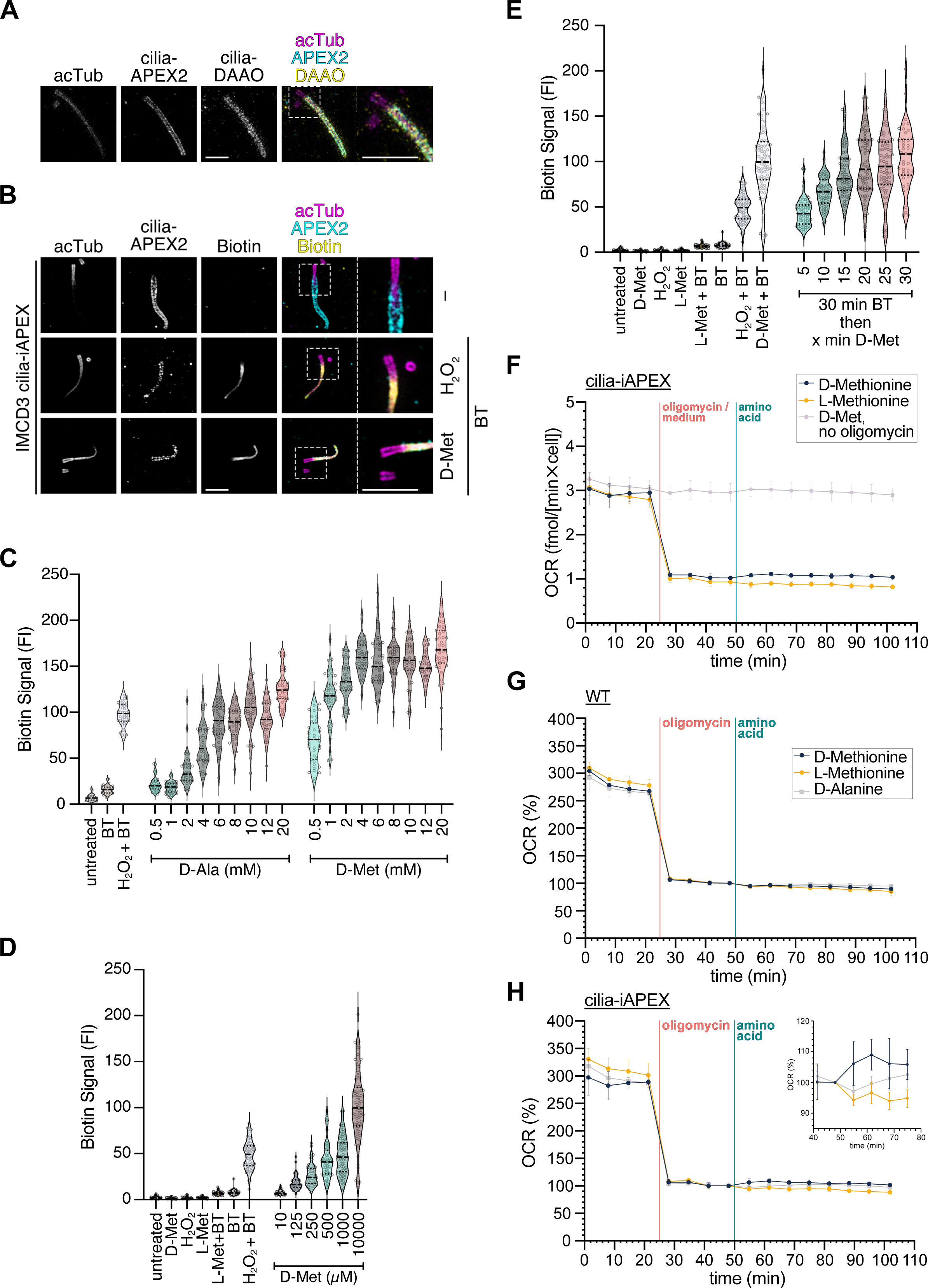
*In situ* D-amino acid oxidase-mediated hydrogen peroxide production enables APEX2 proximity labeling. **(A** and **B)** Ultrastructure Expansion microscopy (U-ExM) confocal images showing cilia-targeted iAPEX localization and proximity labeling in two different cell lines. Cells were fixed, cross-linked, and embedded in a water-expandable gel. After denaturation, they were probed with antibodies against acetylated tubulin, GFP (cilia-APEX2), and ALFA (cilia-DAAO). **(A)** An expanded primary cilium from an RPE-1 cell line stably expressing cilia-iAPEX reveals membrane localization of APEX2 and DAAO. **(B)** U-ExM micrographs of IMCD3 cells stably expressing cilia-iAPEX demonstrates biotinylation within the entire cilium. Cells were either untreated (–), labeled with biotin tyramide and H_2_O_2_ (1 mM, 3 min), or 10 mM D-methionine (D-Met, 30 min). Biotin was visualized using fluorescently labeled streptavidin. Scale bars = 5 µm (adjusted to expansion factors = 4 (D) and 4.3 (E)). **(C**, **D and E)** Quantification of absolute ciliary biotin fluorescence signals in micrographs obtained from proximity labeling experiments performed in IMCD3 cilia-iAPEX cell line shown as violin plots. Quartiles and medians are represented by dotted and dashed lines, respectively. **(C)** Type and concentration of D-amino acid affect biotinylation. Indicated concentrations of D-Ala or D-Met were incubated for 30 min. n = 40 cilia per condition. **(D)** DAAO shows stereoselectivity for D-amino acids and allows labeling with low concentrations of D-Met. n = 77 randomized cilia from two experiments. **(E)** Shorter substrate incubation leads to comparable biotinylation as H_2_O_2_-induced labeling. Cells were incubated with BT and 10 mM D-Met for indicated times. n = 20 randomized cilia from two experiments. Where indicated 1 mM H_2_O_2_ was incubated for 3 min. **(F, G and H)** D-Met-activated cilia-DAAO generates minute amounts of H_2_O_2_. O_2_ consumption rates (OCR) have been measured by Seahorse metabolic flux analysis. **(F)** OCRs of cilia-iAPEX IMCD3 cells were determined after treatment with or without 1.5 µM oligomycin to block cellular respiration, followed by D-Met or L-Met addition to activate DAAO. **(G)** OCRs of wild-type (WT) and **(H)** cilia-iAPEX IMCD3 cells were recorded and normalized to OCR after oligomycin treatment before addition of indicated amino acids (100%). Lines show means of three measurements; error bars depict standard deviations (n = 3).

To identify the minimum concentrations of D-amino acids required for efficient labeling, we titrated the DAAO substrates D-alanine (D-Ala) and D-methionine (D-Met) and assessed cilia-iAPEX-catalyzed biotinylation efficiency by immunofluorescence microscopy (**Fig. 2C** and **Fig. S2A**). Quantitation of biotinylation in primary cilia showed a concentration-dependent increase in biotinylation for both D-Ala and D-Met (**Fig 2C**). High concentrations of both D-amino acids led to stronger biotinylation in cilia compared to 3 min labeling with H_2_O_2_, although D-Ala did not reach the same levels as D-Met. For D-Met saturating signals were achieved at 4 mM (**Fig. 2C**), while notable biotin signals could be observed at concentrations as low as 125 µM when DAAO-catalyzed proximity biotinylation was performed for 30 min (**Fig. 2D**). We therefore focused on the use of D-Met as DAAO substrate for our applications. As biotin tyramide exhibits moderate cell permeability^2^, we aimed to increase the temporal resolution of biotinylation by pre-incubating cells with biotin tyramide before D-Met addition. A time course of the labeling reaction revealed that 5 min incubation with 10 mM D-Met after 30 min biotin tyramide pre-incubation was comparable to short activation (3 min) with 1 mM H_2_O_2_ (**Fig. 2E**), which is compatible with previous time-resolved proteomics applications^17^. Further experiments revealed that 5 min pre-incubation with biotin tyramide was sufficient, as longer pre-incubation times did not increase the labeling efficiency (**Fig. S2B**). Interestingly, pre-incubation with D-Met to initiate local H_2_O_2_ production prior to biotin tyramide addition decreased biotinylation in a time-dependent manner (**Fig. S2B**), indicating that prolonged H_2_O_2_ production might interfere with APEX2 function. Thus, the overall temporal resolution that can be achieved in IMCD3 cells is 5 min, which compares favorably to recently developed proximity labeling methods.

To assess potential toxicity of the iAPEX system, we determined the H_2_O_2_ production by cilia-DAAO employing oxygen (O_2_) consumption measurements as DAAO activity converts O_2_ to equimolar amounts of H_2_O_2_^37^. When blocking cellular respiration with oligomycin the cilia-iAPEX IMCD3 cell line consumed approximately 1 fmol/(min×cell) O_2_ at steady state (**Fig. 2F**). After addition of D-Met the O_2_ consumption increased to about 1.2 fmol/(min×cell) (**Fig. 2F**). This rise in O_2_ consumption was D-amino-acid-and cilia-DAAO-dependent, as it was not observed in a parental cell line (**Fig. 2G**). Although D-Ala led to a comparable maximum O_2_ consumption rate as D-Met, our kinetic analysis indicated a 30 min delay to reach this maximum (**Fig. 2H**), which agrees with the observed differences in iAPEX-catalyzed biotinylation (see **Fig. 2C**). Assuming an average cell volume of 4000 fL, the increase in O_2_ consumption of 0.2 fmol/(min×cell) would result in sub-µM H_2_O_2_ concentrations within 1 sec of DAAO activity if the cell completely lacked mechanisms to detoxify H_2_O_2_. However, as physiological redox signaling requires an existing, potent antioxidant system^38^, our data indicate that the amount of H_2_O_2_ produced by cilia-DAAO is within physiological H_2_O_2_ concentrations and can therefore be considered non-toxic for most cell types.

### Cilia-iAPEX locally restricts proximity biotinylation and prevents off-target biotinylation

To test whether the iAPEX labeling cascade overcomes the high background observed in select cell lines (**see Fig. 1A,B**), we introduced cilia-iAPEX into NIH/3T3 cells using FlpIn recombinase, and isolated a clonal cell line, in which both enzymes localized to primary cilia (**Fig. 3A**). Most importantly, iAPEX labeling was specific to primary cilia in this cell line and indicated that spatially restricted H_2_O_2_ production by cilia-DAAO prevented non-specific biotinylation of other cellular structures, as observed after direct activation of cellular peroxidases (**Fig. 3A**). SDS-PAGE and Western Blot analysis of cilia-iAPEX expressing NIH/3T3 cells confirmed high background biotinylation when using H_2_O_2_ as a substrate, while DAAO activation resulted in markedly reduced but specific biotinylation (**Fig. 3B**). Interestingly, even in the established IMCD3 cell line we observed overall stronger signals when using H_2_O_2_ compared to D-Met-based proximity labeling (**Fig. 3B**, lanes 8 *vs.* 9), despite weaker signals in primary cilia (see **Fig. 2C**). To gain a deeper understanding of the events occurring during APEX labeling we established a live-cell imaging setup to visualize the subcellular localization of peroxidase activity by the oxidation of the peroxidase substrate Amplex UltraRed (AmUR) to a fluorescent resorufin product^39,40^. After loading cells that stably express cilia-iAPEX with AmUR we noticed a burst in peroxidase activity throughout the entire cell shortly after H_2_O_2_ addition, which ceased over time when only the APEX activity in the primary cilium remained (**Fig. 3C** and **Video 1**). In contrast, local production of H_2_O_2_ by cilia-DAAO prevented off-target peroxidase activity, as we observed resorufin signals exclusively in primary cilia for prolonged labeling times (**Fig. 3D** and **Video 2**). These results indicated that other cellular peroxidases are capable of oxidizing substrates, such as biotin tyramide to biotinylate nearby proteins by proximity labeling when H_2_O_2_ is added to the cells. We further hypothesized that the observed initial burst in peroxidase activity may cause significant off-target biotinylation and thereby high background when studying proteomic environments of targets that are expressed at low levels, such as for primary cilia proteomics.

**Fig. 3:**
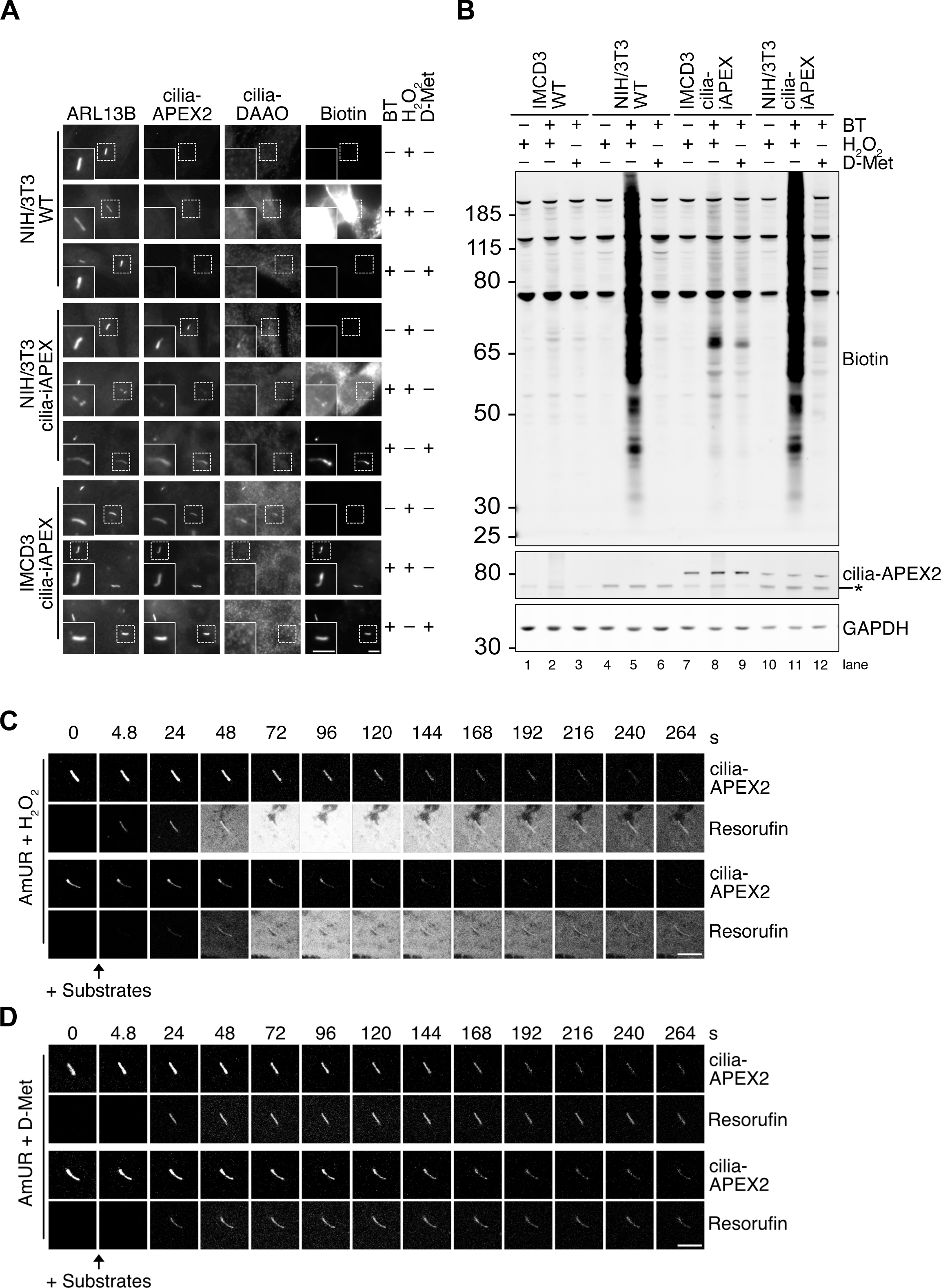
iAPEX enables cilia-specific biotinylation in NIH/3T3 bypassing high cellular background. **(A)** IF micrographs of NIH/3T3 WT and cilia-iAPEX cells with IMCD3 cilia-iAPEX cells as control. Cells were left untreated, labeled using BT and H_2_O_2_, or BT and D-Met, as indicated. **(B)** Western blot analysis of WT and cilia-iAPEX-expressing IMCD3 and NIH/3T3 cells. Cells were lysed before or after BT and H_2_O_2_, or BT and D-Met treatment. Asterisk marks cross-reactive band of the anti-GFP antibody. **(C and D)** Live-cell confocal imaging micrographs were captured to observe peroxidase-dependent Amplex UltraRed (AmUR) oxidation to resorufin in IMCD3 cells stably expressing cilia-APEX2. **(C)** Cells were treated with 50 µM AmUR together with 1 mM H_2_O_2_ where indicated. Resorufin and GFP autofluorescence of cilia-APEX2 were monitored at 4.8-second intervals over a total duration of 264 s (see also **Video 1**). **(D)** AmUR oxidation reveals exclusive cilia-APEX2 activity in DAAO-dependent proximity labeling after addition of 50 µM AmUR and 10 mM D-Met (see also **Video 2**). Scale bars = 5 µm in all panels.

### cilia-iAPEX proteomics increases specificity and sensitivity of cilia protein identification

To directly compare the iAPEX-with the APEX2-based proximity labeling method as a discovery tool in proteomics applications, we performed iAPEX (DAAO-dependent) and APEX2 (H_2_O_2_) proximity labeling in cilia-iAPEX IMCD3 cells using desthiobiotin tyramide (DTBT) as a substrate, as this allowed competitive elution of APEX-biotinylated proteins after isolation by streptavidin chromatography (**Fig 4A**). Abundant non-specific biotinylation in APEX2-based proximity labeling setups requires controls to precisely assess the background^41,42^. To this end, for cilia proteomics we previously expressed cilia-APEX2 in *Cep164^-/-^* cells that lack primary cilia^17^. Triplicates of the iAPEX-labeled samples and duplicates of the controls were analyzed by SDS-PAGE and Western blotting. Our analyses confirmed reduced biotinylation by iAPEX compared to APEX2 labeling (**Fig. S3A**, lanes 7-8 vs. 9-11), while several cilia components, represented by IFT88 and IFT57, were isolated more efficiently after iAPEX labeling (**Fig. 4B**, lanes 13-15 *vs.* 11-12). This indicated a higher sensitivity of cilia-iAPEX compared to previous setups, while the *Cep164^-/-^* controls confirmed specificity of isolation, as no ciliary proteins were isolated in the absence of cilia (**Fig. 4B**, lanes 9-10). To quantitatively assess the performance of cilia-iAPEX vs. cilia-APEX2, isolated proteins were digested by trypsin, labeled with tandem-mass-tags (TMT) and analyzed by synchronous precursor selection mass spectrometry (SPS-MS^3^) (**Fig. 4A**;^5,17,43,44^). We quantified the relative abundances of 5982 identified proteins within the individual samples (see **Table S1**). When assessing candidate ciliary proteins by statistical analysis of relative enrichments between cilia-iAPEX and control samples we applied stringent TMT enrichment ratios of 2^3^ which resulted in 175 high confidence candidate cilia proteins (**Fig. S3B**). Surprisingly, within the same experiment the same TMT enrichment ratio cutoff between cilia-APEX2 and control samples identified 799 putative cilia proteins (**Fig. S3C**). A direct comparison showed that the enrichment of known cilia proteins was similar in both approaches, however, cilia-iAPEX proteomics separated known cilia proteins much better from false-positives (**Fig. 4C**). Gene Ontology (GO) term enrichment analyses confirmed higher specificity of the iAPEX setup, as evidenced by the absence of non-ciliary processes and the lower *p* values of ciliary categories (**Fig. S3D,E**). Hierarchical clustering of the relative protein abundances within the experiment demonstrated high reproducibility of the replicate samples (**Fig. S4A**). Proteins enriched in both iAPEX and APEX2 labeled samples formed three clusters highly enriched in known cilia proteins (**Figs. 4D**, **S4B** and **Table S1**), which covered 38% of the cilia-APEX2 proteome^17^. A GO_term enrichment analysis revealed high statistical significance of components associated with cilia and related microtubule-based structures (**Fig. 4E**), while our previous cilia-APEX2 proteome contained many non-ciliary categories^17^, suggesting false-positive hits. Agreeingly, our cluster analysis also identified proteins that were enriched only in the H_2_O_2_-treated samples, in wild-type and *Cep164^-/-^* cells, that formed four clusters (**Figs. 4F** and **S4C**). GO_term enrichment analysis of these clusters identified a large fraction of proteins located in the endoplasmic reticulum (ER) (**Fig. 4G**), suggesting that ER resident peroxidases in IMCD3 cells can biotinylate nearby proteins in a H_2_O_2_-dependent manner. Such peroxidases may cause non-specific labeling and contribute to potential false-positive hits in classic APEX2 labeling setups. Taken together, iAPEX-based proteomics shows high sensitivity to analyze the proteome of subcellular microdomains and significantly reduces the number of false-positives by lowering background biotinylation activities.

**Fig. 4:**
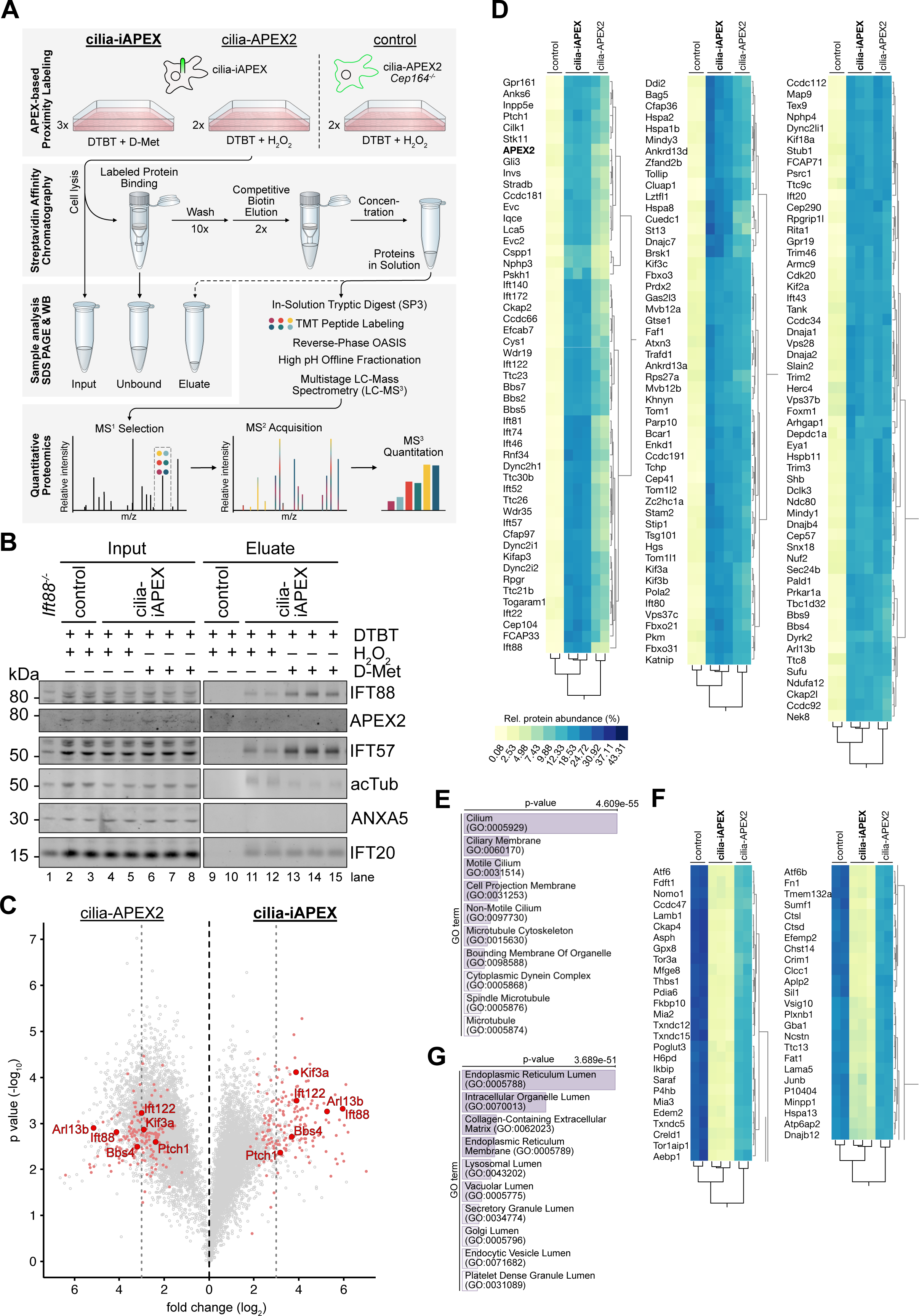
Quantitative primary cilia proteomics using iAPEX outperforms conventional APEX2-based system in IMCD3 cells. **(A)** Schematic illustration of cilia-iAPEX-based proximity labeling workflow for proteomic analysis of IMCD3 primary cilia. cilia-iAPEX labeling was performed in cells expressing cilia-iAPEX (cilia-APEX2 and cilia-DAAO) with desthiobiotin tyramide (DTBT) and D-Met for 30 min. For cilia-APEX2 proximity labeling cilia-iAPEX cells were pre-incubated with DTBT for 30 min, followed by 3 min H_2_O_2_, which was also performed in cilia-ablated *Cep164^-/-^* cells expressing cilia-APEX2 as controls. After labeling, cells were lysed, labeled proteins isolated by streptavidin affinity chromatography and competitively eluted with biotin. Input, Unbound and Eluate fractions can be analyzed by SDS-PAGE and Western Blotting. For mass spectrometric analysis, eluted proteins were digested in-solution using trypsin, peptides labeled with TMTs (tandem mass tags) and fractionated offline via reverse-phase OASIS chromatography. Quantitative proteomics was performed using LC-MS³, where peptides were selected (MS¹), fragmented for identification (MS²), and TMT reporter ions quantified (MS³). (B) Western blot analysis of samples after proximity labeling from IMCD3 cilia-iAPEX or cilia-ablated cilia-APEX2 *Cep164^-/-^*cells (control), as outlined in **(A)**. Lysate from IMCD3 *Ift88^-/-^*cells served as an antibody specificity and untreated control. Input and Eluate samples were separated by SDS-PAGE and analyzed by Western Blot using indicated antibodies. Increased amounts of cilia proteins, represented by IFT88 and IFT57, were isolated after DAAO-dependent proximity labeling. Input 0.063 %, Eluate 8.5 %. (C) Volcano plot of statistical significance *versus* protein enrichment in cilia-APEX2 (left) and cilia-iAPEX (right) compared with control samples. Calculated *p* values (unpaired Student’s *t* test) were plotted against TMT ratios for 5982 proteins. Proteins are indicated by grey circles; red circles show known cilia proteins. Representative subunits of kinesin-2 (Kif3a), IFT-A (Ift122), IFT-B (Ift88) and the BBSome (Bbs4) are highlighted. Dotted lines indicate TMT ratios of 2^3^. (D) Selected clusters of two-way hierarchical cluster analysis of IMCD3 cilia-iAPEX proteome show known cilia proteins and highest scoring candidate cilia proteins. Legend shows relative protein abundances (in %). Full cluster shown in **Fig. S4**. (E) GO_term enrichment analysis of protein clusters in (D) shows enrichment of non-ciliary categories in samples treated with H_2_O_2_. *p* values were calculated by Fisher’s exact test. (F) Selected clusters with proteins identified after H_2_O_2_-mediated cilia-APEX2 proximity labeling. (G) GO_term enrichment analysis of protein clusters in (F) identified enrichment of non-ciliary categories in samples treated with H_2_O_2_. *p* values were calculated by Fisher’s exact test.

### Determining the cilia-iAPEX proteome of NIH/3T3 cells

As iAPEX proximity labeling allowed the specific biotinylation of proteins in NIH/3T3 cells (see **Fig. 3**), we sought to gain proteomic information on the ill-defined proteome of NIH/3T3 primary cilia. By combining hierarchical clustering with the increased specificity of the iAPEX system, we envisioned that the cilia proteomes could be investigated without the need for genetic background controls, such as the *Cep164^-/-^*. Therefore, we utilized the cilia-iAPEX NIH/3T3 cell line and included two simple specificity controls that prevent iAPEX labeling: 1) omitting the APEX substrate desthiobiotin tyramide, and 2) replacing the DAAO substrate D-Met for L-Met which does not activate DAAO (**Fig. 5A**, see also **Figs. 2D,E**). Replicate samples were prepared, followed by streptavidin chromatography and analysis by SDS-PAGE and Western Blotting. The analysis revealed biotinylation of proteins in the presence of desthiobiotin tyramide (**Fig. 5B**), independent of DAAO activation, which suggests background labeling mediated by endogenous H_2_O_2_. Nonetheless, we observed an increase in biotinylation when DAAO was activated by D-Met (compare lanes 5-7 to 11-13). Importantly, there was a clear enrichment of the cilia proteins IFT57 and IFT88 only when iAPEX labeling was performed (**Fig. 5C**), which indicated that cilia proteins were only labeled when both enzymes were activated and omitting either substrate could serve as specificity controls. We therefore continued with TMT labeling and SPS-MS^3^ analysis, followed by hierarchical clustering of the relative abundances of the 6067 quantified proteins (see **Fig. S5A** and **Table S2**). Most clusters contained similarly distributed protein abundances, indicating non-specific background binding. In 14 clusters, proteins were enriched in the six desthiobiotin tyramide-treated samples, indicating biotinylation (see also **Fig. 5B**). Only four out of those 14 clusters contained proteins highly enriched only in the iAPEX-labeled samples and included many known cilia proteins, such as IFT subunits, BBSome components and molecular motors (**Figs. 5D**, **S5B** and **Table S2**). These clusters also contained several proteins previously not implicated in cilia biology. When comparing the cilia-iAPEX proteomes of IMCD3 cells with NIH/3T3 cells, we noticed an overlap of below 50% (**Fig. 5E** and **Table S3**), which could suggest cell type-specific heterogeneity of the cilia proteome. Both datasets revealed novel cilia candidate proteins. We therefore set out to confirm cilia localization of select candidates by independent methods. To this end we tagged the proteins of interest with an ALFA affinity tag^45^, transiently transfected IMCD3 cells and assessed the localization of the respective proteins by fluorescence microscopy (**Fig. 5F**). Indeed, we could confirm the localization of the proteins PSKH1, CUEDC1, and CKAP2L to the primary cilium shaft. We could also validate cilia localization of two so-far uncharacterized mouse homologs of the human open reading frames C19orf44 and C7orf57 in IMCD3 cells, which we termed FCAP71 and FCAP33 (Found in cilia-iAPEX proteome of 71 and 33 kDa), respectively. FCAP33 showed enrichment in primary cilia, whereas FCAP71 localized to the base of cilia, marked by CEP164 (**Fig. 5F**). While the functions of these novel cilia proteins remain to be investigated, we conclude that iAPEX proximity labeling is a powerful, unbiased discovery tool for subcellular proteomics of previously inaccessible cell lines.

**Fig. 5:**
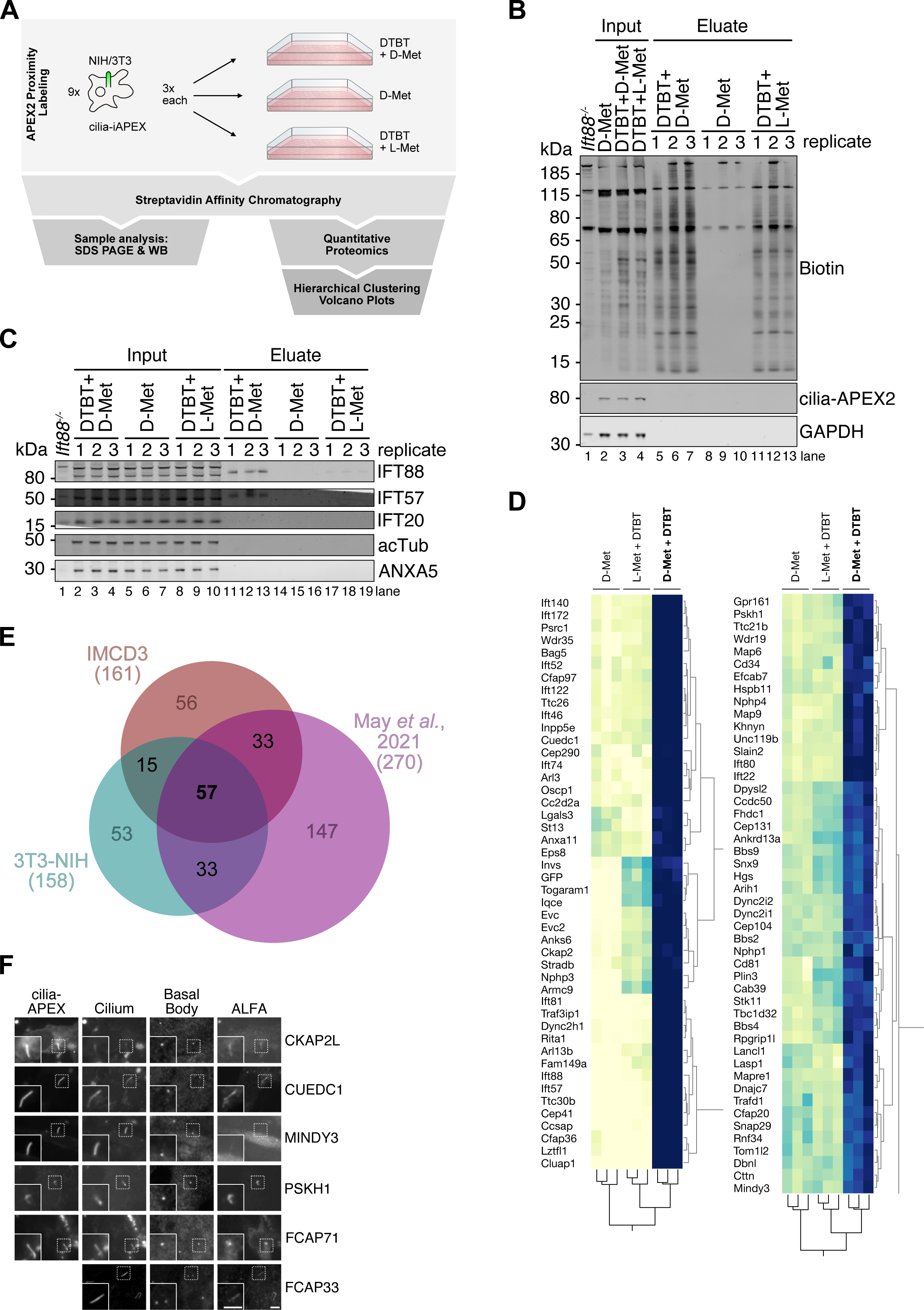
iAPEX allows specific quantitative primary cilia mapping in previously for APEX2 inaccessible NIH/3T3 cells. **(A)** Scheme of cilia-iAPEX proximity labeling workflow for primary cilia proteomics in NIH/3T3 cells. DAAO-dependent proximity labeling was performed by incubating cells with desthiobiotin tyramide (DTBT) and D-Met. As controls, cells were incubated with DTBT and L-Met, or D-Met only. Sample processing according to schematic in Fig. 4A. **(B and C)** After proximity labeling, cells were lysed, biotinylated proteins enriched by streptavidin chromatography, and Input and Eluate samples analyzed by SDS-PAGE and Western blotting analysis. **(B)** DTBT incubation causes background biotinylation. Biotin was detected using fluorescently labeled streptavidin, cilia-APEX2 by GFP-specific antibodies. Input 0.063 %, Eluate 1.2 %. **(C)** Cilia proteins were specifically isolated after cilia-iAPEX labeling. Indicated proteins were detected using specific antibodies. IMCD3 *Ift88^-/-^* cells served as an antibody control. Input 0.063 %, Eluate 8.8 %. **(D)** Two-way hierarchical cluster analysis of NIH/3T3 cell cilia-iAPEX proteome. Zoom on clusters highly enriched in cilia proteins. Full cluster shown in **Fig. S5**. **(E)** cilia-iAPEX proteomes of IMCD3 and NIH/3T3 cells show distinct overlap with cell specific differences. Venn diagram depicting proteomic overlap of iAPEX-proximity labeled IMCD3 and NIH/3T3 cells against the cilia-APEX2 proteome^17^. **(F)** Validation of primary cilia localization of proteins previously not linked to cilia. The representative IF micrographs illustrate IMCD3 cells transiently transfected with plasmids expressing both cilia-APEX2 (as transfection and localization control) and indicated primary cilia candidate proteins C-terminally fused to an ALFA-tag or ALFA-tag fusion alone (for FCAP33). Upon fixation, primary cilia and basal bodies were visualized using antibodies targeting ARL13B or acTub (Cilium) and CEP164 or γTub (Basal Body) respectively, while the proteins of interest (POI) were stained with an anti-ALFA antibody. Scale bars = 5 µm.

### Expanding the iAPEX enzyme cascade to other cell types and organisms for *in situ* applications

For cilia proteomics in IMCD3 and NIH/3T3 cells, the iAPEX transgenes were integrated into existing FRT sites on the genomes of the respective cell lines. For a more versatile delivery of the transgenes into additional cell lines of interest, such as C2C12 myoblasts, 3T3-L1 pre-adipocytes or primary cells, we engineered plasmids for packaging NPHP3^1–200^–EGFP-APEX2 (cilia-APEX2) and NPHP3^1–200^–FLAG-DAAO-ALFA (cilia-DAAO) transgenes separated by an internal ribosome entry site (IRES) into lentiviral particles for cell infection (**Fig. 6A**) After lentivirus infection, GFP-positive cells were sorted by fluorescence-activated cell sorting (FACS) and subjected to H_2_O_2_-based APEX2 labeling or iAPEX labeling using D-Met. Analysis by fluorescence microscopy revealed that both cilia-APEX2 and cilia-DAAO exhibited specific localization to primary cilia in the respective cell types (**Fig. 6B**). Despite the specific localization of both enzymes, exogenous H_2_O_2_-based APEX2 proximity labeling resulted in significant background biotinylation, which explains previous limitations in performing cilia proteomics with these cell lines (K. Hilgendorf and D. Mick, personal communication). In contrast, iAPEX labeling using D-Met led to specific biotinylation within primary cilia with strongly reduced background in our fluorescence microscopy setup. SDS-PAGE and Western Blot analysis confirmed immense background biotinylation when using H_2_O_2_ (**Fig. 6C**, lanes 13-16), while D-Met-based labeling reduced the biotinylation to levels observed in the cilia-iAPEX IMCD3 cell line (**Fig. 6C**, lanes 9-12). While determining the primary cilia proteomes of both C2C12 and 3T3-L1 cell types brings additional challenges due to the relatively low ciliation rates^46–50^ and the resulting low amounts of labeled cilia proteins, we are convinced that our improved iAPEX workflow will finally enable mass spectrometry-based characterization of the cilia proteome during dynamic cilia processes, such as cell differentiation.

**Fig. 6:**
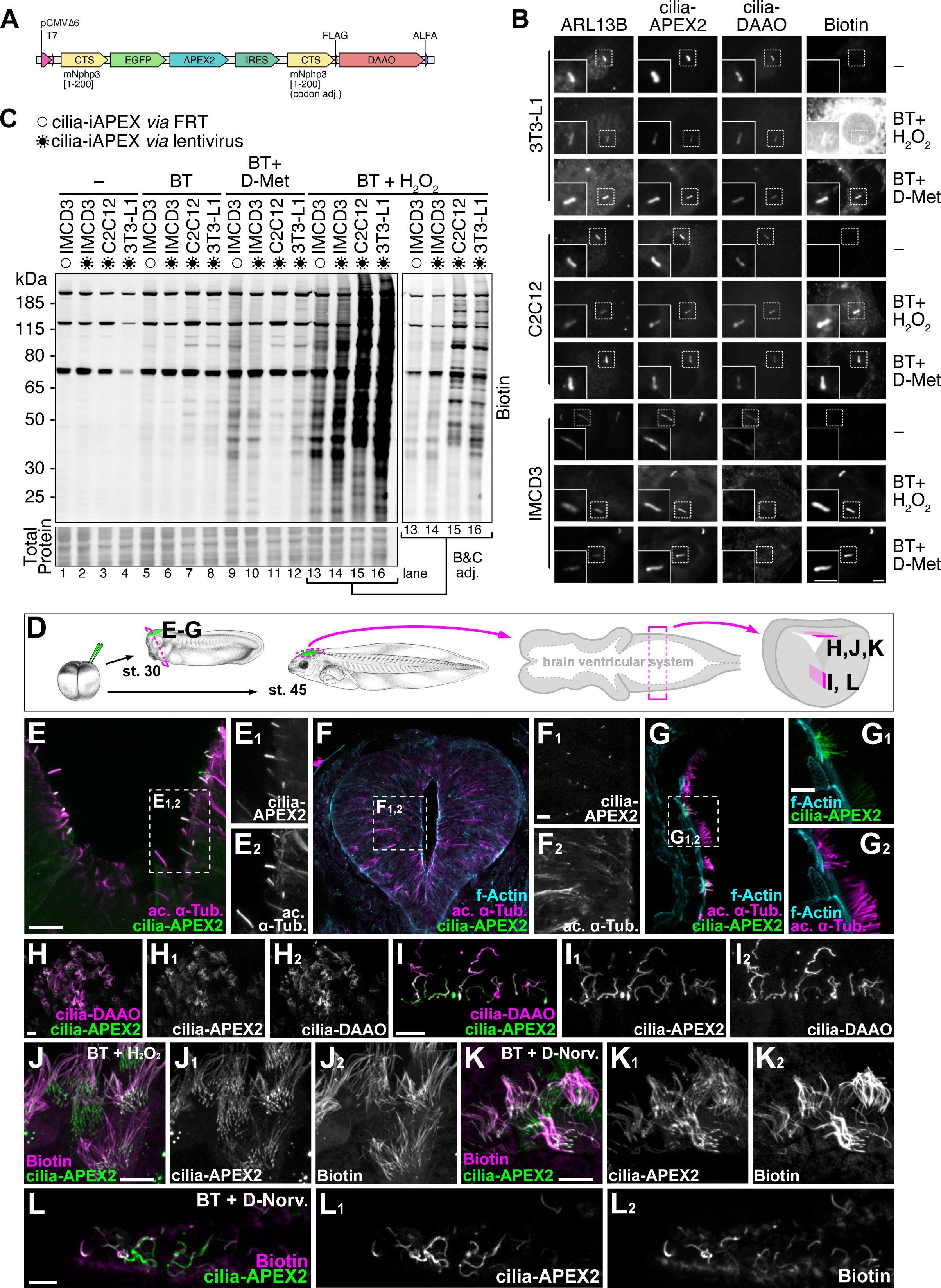
Establishing iAPEX-based proximity labeling in previously inaccessible cell types and *Xenopus laevis*. **(A)** Engineered lentiviral transfer vector harboring a polycistronic cassette for the cilia-iAPEX two-component expression. Transcription is controlled by low-expressing truncated CMV promoter (P_CMVΔ6_). cilia-APEX2 and cilia-DAAO transgenes are separated by an internal ribosome entry site (IRES). Greyed out elements are common transfer vector components needed in second generation lentiviral systems. **(B)** 3T3-L1, C2C12 and IMCD3 cell lines have been infected with lentiviral vectors to express the cilia-iAPEX transgenes (depicted in (A)). Following proximity labeling with biotin tyramide (BT) and H_2_O_2_ or D-Met as indicated, cells were fixed and processed for immunofluorescence microscopy using antibodies specific to ARL13B to mark cilia, and ALFA tag to detect cilia-DAAO. Biotinylation was visualized using fluorescently labeled streptavidin and cilia-APEX2 by GFP fluorescence. Scale bars = 5 µm. **(C)** After incubation with indicated reagents for proximity labeling, IMCD3, C2C12, and 3T3-L1 cells expressing cilia-iAPEX after lentivirus infection (virus symbols) were lyzed and analyzed by SDS-PAGE and Western blotting. cilia-iAPEX IMCD3 cells, in which the transgenes are expressed from the FlpIn locus served as control (empty circles). Biotin was visualized using fluorescently labeled streptavidin, equal protein loading (25 µg/lane) confirmed by total protein stain. Lanes 13-16 were brightness and contrast adjusted to visualize banding patterns. Where indicated BT and D-Met have been incubated for 30 min, H_2_O_2_ for 3 min. **(D)** mRNA transcribed from plasmids containing cilia-APEX2 or cilia-iAPEX cassettes was injected into one or two dorsal animal blastomeres of four-to-eight-cell *Xenopus laevis* embryos to target constructs to the central nervous system. After rearing to tailbud / tadpole stages (st. 30 / 45), hemisections through the brain area (st. 30) or brain preparations (st. 45) were immunostained to visualize enzyme expression using anti-GFP and anti-ALFA-tag immunostaining for cilia-APEX2 and cilia-DAAO, respectively. **(E-G)** APEX2, expressed from cilia-APEX2 (E) or cilia-iAPEX (F, G) constructs, localizes to acetylated α-Tubulin (ac. α-Tub.)-positive primary cilia of the neural tube floor plate (E) and lateral neural tube (F) and to cilia of multiciliated epidermal cells (G). Scale bars: 10 µm. **(H, I)** Co-localization of cilia-APEX2 and cilia-DAAO expressed from cilia-iAPEX constructs in multiciliated cells of the roof (H) and floor plate (I) in the tadpole hindbrain. Scale bars: 10 µm. **(J-L)** Biotinylation using cilia-iAPEX, fluorescently labeled strepavidin detects biotin in st. 45 hindbrain roof multiciliated cells (J, K) and floorplate monociliated cells (L). (J) Biotin tyramide (BT) and H_2_O_2_ were sequentially injected into the hindbrain ventricle of st. 45 tadpoles *in vivo*, followed by fixation of whole embryos, brain dissection and (immuno)staining. (K, J) After 20 min fixation of whole embryos, brains were dissected and incubated in BT and D-Norvaline for 3 min, followed by quenching and (immuno)staining. Scale bars: 10 µm.

To further demonstrate the versatility of the iAPEX system, we expressed both transgenes *in vivo* in the developing *Xenopus laevis* embryo by RNA injections (**Fig. 6D**). At early embryonic development we found high enrichment of cilia-APEX2 in cilia of diverse tissues, such as primary cilia in the neural tube (**Figs. 6E-F**) and the multiciliated cells of the epidermis (**Fig. 6G**). To assess the functionality of the iAPEX enzymatic cascade in complex tissues, we investigated the brain ventricular system at later embryonic stages and confirmed specific co-localization of both cilia-APEX2 and cilia-DAAO to cilia of multiciliated cells of the hindbrain roof (**Fig. 6H**) and monociliated cells of the floorplate (**Fig. 6I**). Injection of the APEX substrates into the ventricular system led to robust biotinylation, as evidenced by fluorescence microscopy of multiciliated hindbrain roof ependymal cells (**Fig. 6J**). Moreover, cilia-iAPEX labeling was achieved in multiciliated (**Fig. 6K**) and monociliated cells (**Fig. 6L**) of dissected brains post-fixation, which confirms the suitability of the new iAPEX system for future *in situ* and *ex vivo* applications.

## DISCUSSION

Proximity labeling technologies have emerged as powerful tools, especially for subcellular proteomics. The principle of an enzymatic activity that labels targets in proximity brings three major advantages: 1) structures that can be marked by transgenes but cannot be purified by other means become accessible to proteomic investigation; 2) proximity labeling will capture low affinity and transient interactions^51^; 3) labeling with enzymatic activities allows for a high temporal resolution of proteomic analysis^2,22^. The latter is a particular strong feature of ascorbate peroxidase (APEX), as its high enzymatic activity allows for sub-minute labeling times, which cannot be achieved by any other available methodology^52^. Yet, as the APEX activity requires hydrogen peroxide, there had been limitations for a more general use, as the toxicity of externally added H_2_O_2_ limits its use in *in vivo* applications^53–55^. Moreover, the presence of endogenous peroxidases generates varying degrees of background signals that precludes specific labeling in many models -especially when studying structures of low abundance, such as the primary cilium (see **Figs. 1A-B**). The iAPEX (improved APEX) enzymatic cascade solves this problem by locally producing H_2_O_2_ and thereby suppresses cell toxicity (see **Fig. 2**) and increases labeling specificity (see **Fig. 4**). While other proximity labeling methodologies remain powerful alternatives with individual strengths and weaknesses, the iAPEX technology still allows a very high temporal resolution of labeling, due to fast enzyme kinetics and the use of substrates that are bio-orthogonal (biotin tyramide) or very rare (D-amino acids) in most biological systems^56^. Importantly, the system is simple to use, as we can deliver both transgenes in one vector (see **Figs. 1C** and **6A**), and the required substrates are commercially available at low costs.

Central to the iAPEX technology is the D-amino acid oxidase (DAAO) from *Rhodotorula gracilis*, which has been used in a wide variety of *in vivo* and *ex vivo* applications^34,57,58^. DAAO oxidizes a broad spectrum of D-amino acids in an FAD-dependent manner. Re-oxidation of FAD results in reduction of molecular oxygen (O_2_) to H_2_O_2_ -the desired product-that is rapidly reduced by cellular peroxins, while the oxidized imino acids are metabolized to keto-acids and ammonium^29^. This molecular mechanism also points at potential limitations of the iAPEX system, which requires molecular oxygen and might cause metabolic imbalances. However, most tissues harbor much higher keto acid and NH_3_ concentrations than DAAO-generated H_2_O_2_^58,59^. In the context of a cellular substructure, such as the primary cilium, we have noticed very low levels of H_2_O_2_ production (see **Fig. 2**) with minimal impact on biological processes in the final steps of sample preparation. O_2_ availability is expected to affect DAAO activity, particularly when investigating tissues with low oxygenation in *in vivo* applications. Optimized DAAO variants, such as mDAAO that shows high activity at lower O_2_ and amino acid concentrations, may solve these issues^60,61^. Indeed, the more relevant limitation of the system we see is cellular availability of D-amino acids, which have to traverse the plasma membrane *via* amino acid transporters^37,38,61–63^. Therefore, cell type-specific differences can be expected, and D-amino acid concentrations and incubation times likely require optimization. Similar limitations in the uptake of substrates for proximity labeling should be considered for all available technologies^2,64^.

### iAPEX has the potential to bypass more complex genetic controls

Our study demonstrates that due to varying levels of endogenous peroxidases, cell types will show different background profiles, which can have a detrimental impact for proteomics of small subcellular structures. Our data also show that even without H_2_O_2_ generation by DAAO, prolonged incubation with tyramides results in biotinylation, which can be attributed to varying levels of endogenous H_2_O_2_. Such background can be revealed by genetic controls, such as cell lines that express mis-localized enzymes (see **Fig. S2A**) or lack the entire structure of interest –such as cilia-less *Cep164^-/-^* cells. However, generating genetic control cell lines is also flawed by potential genetic drift leading to proteomic alterations^65–67^. While isogenic control cell lines might still be the gold standard, we could show that a single iAPEX cell line can be employed to determine the cilia proteome of NIH/3T3 cells. Including information from samples with and without the H_2_O_2_ production by DAAO improved clustering resolution and circumvented the need for additional genetic controls - a major advantage for future *in vivo* applications.

In this proof-of-concept study we utilized the increase in sensitivity and specificity of the iAPEX technology to study the primary cilium proteome. A systematic comparison allowed us to identify several false-positive candidate cilia proteins from our own and other previous studies^13,14,17^, such as proteins with known function in the endoplasmic reticulum or mitochondria. Despite a common realization that many proteins exist at multiple subcellular locations^68^, it appeared unlikely that all previous hits fulfilled additional functions in primary cilia, and we now provide experimental evidence that they represented unspecific background in previous studies.

### Comparative cilia proteomics

By omitting H_2_O_2_ addition, iAPEX allows for much longer labeling times than with externally added H_2_O_2_, which not only reduces toxicity but also off-target labeling effects at extended labeling times. Thereby, iAPEX labeling leads to stronger biotinylation and increased sensitivity. This allowed us, despite the much smaller scale compared to previous studies^13,17^, to identify new candidate cilia proteins with unknown functions in cilia biology, not only in the well-studied IMCD3 model but also in previously inaccessible NIH/3T3 primary cilia, which had only been investigated using TurboID^18^. Here, we revealed an overlap of only about 45% between the cilia proteomes of these two cell lines. While the observed differences may support previous suggestions of cilia heterogeneity, they may also be caused by experimental variation in single mass spectrometry runs^69^. Nonetheless, our proof-of-concept study confirms that core cilia proteins, such as IFT, BBSome, kinesin and dynein subunits seem to be common to primary cilia of both cell types, while we identified many signaling components, such as kinases and putative transcription factors that differed between cell lines, which might explain clinical differences observed in ciliopathies^7,70,71^. With the iAPEX system available and the ease of use, we anticipate that many more cell type-specific cilia proteomes will become available soon to tackle this larger question in the cilia community.

At the same time, we also want to highlight the primary cilium as an interesting model to study enzyme reactions in a defined cellular microdomain. By using cilia targeting signals, we directed enzymes into the primary cilium and reconstituted a DAAO-APEX enzymatic cascade, which can be studied in detail (substrate and product concentrations, reaction times and temperatures etc.) in a unique *in cellulo* environment. The primary cilium may therefore also be recognized as a “living test tube” with specific geometry and unique biophysical properties that may be a powerful model for future synthetic biology applications.

### Comparison to similar technologies

While we used the primary cilium as an example for a cellular microdomain that is difficult to purify, the iAPEX system promises to be applicable to other cellular substructures^72,73^. The iAPEX system requires two enzymatic activities to co-operate, which opens possibilities for proximity labeling of subpopulations of proteins or structures that are defined by the co-localization of two markers. The use of different targeting signals for APEX and DAAO has the potential to increase spatial specificity to probe for subpopulations of organelles, proteins in complex with specific interaction partners, or organelle contact or biogenesis sites^72,73^. Similar co-localization-based experimental setups exist in the forms of split-APEX, where the enzymatic APEX2 activity is reconstituted by complementation of two protein parts^74^, and TransitID, which combines TurboID-with APEX2-based proximity labeling, followed by two-step purification schemes^75^. While the former requires two tagged proteins to directly interact in an orientation that allows APEX2 reconstitution, which may induce non-physiological protein interactions, TransitID is limited by the relatively slow enzymatic kinetics of TurboID labeling. Moreover, both systems require exogenous addition of H_2_O_2_. Therefore, despite clear advantages, split-APEX and TransitID may not be suitable for biological systems with higher levels of endogenous peroxidases that increase background labeling (see **Fig. 1A**), or where biotin scavenging or H_2_O_2_ toxicity should be avoided^76,77^. Since both technologies rely on APEX2 activities, they could -in theory-be “improved” and combined with *in situ* generation of H_2_O_2_ by DAAO or similar enzymes. It remains to be seen whether these methods will be combined by many non-specialists, as this would further increase the complexity of experimental setups. In this regard, we believe that the improved APEX methodology provides a powerful improvement of an existing technology that is simple to implement and greatly increases sensitivity and specificity for subcellular proteomics applications.

## MATERIALS AND METHODS

### Cell culture and cell line generation

Wild-type cells and established cell lines, including 3T3-L1 preadipocytes, C2C12 myoblasts, HEK293T cells, and NIH/3T3 fibroblasts, were cultured in DMEM (Fisher Scientific, Cat. No. 11594486). IMCD3 cells were grown in DMEM/F12 (Fisher Scientific, Cat. No. 11594426). All media were supplemented with 7.5% FBS (Fisher Scientific, Cat. No. 11573397). All cells were propagated at 37 °C, 5% CO_2_. To induce ciliation, cells were serum-deprived in 0.2% FBS containing media for 24 h. IMCD3 cell lines stably expressing cilia-APEX2 and control-APEX2 have been previously described^17^. IMCD3 cell lines stably expressing cilia-iAPEX, cyto-iAPEX, and cilia-DAAO-APEX2, as well as NIH/3T3 cell line stably expressing cilia-iAPEX, were generated using the Flp-In system as previously described^78^. A Flp-In-compatible vector (pEF5B-FRT-cilia-APEX2/cilia-DAAO) with back-to-back CMVΔ6 and EF1α-TATA-box-mutant promoters was generated to enable simultaneous expression of genetically targeted APEX2 and DAAO transgenes. For cloning, DAAO was amplified from pAAV-SypHer2-DAAO-NES (gift from L. Prates Roma). Forward transfections were performed using jetPRIME transfection reagent (VWR, Cat. No. 101000046) according to manufacturer’s guidelines. Cloning cylinders (Sigma-Aldrich, Cat. No. CLS31668) were employed to obtain cell clones.

For 3T3-L1 and C2C12 cells, the multicistronic lentiviral vector pLVX-cilia-APEX2-IRES-cilia-DAAO was designed, which contained a CMVΔ6 promoter and an internal ribosomal entry site (IRES) to enable co-expression of the transgenes cilia-APEX2 and cilia-DAAO. The multicistronic plasmid was synthesized by BioCat. Lentivirus was produced by transfecting HEK293T cells with second-generation lentiviral vectors (psPAX2, pMD2g-VSV-G, and pLVX-cilia-APEX2-IRES-cilia-DAAO in a 1:1:2 ratio). After 24 h, medium was replaced. 24 h later lentivirus containing supernatant was collected, filtered through 0.45 µg PES filter, and supplemented with 4 µg/ml polybrene for infection of IMCD3 and 3T3-L1 and 10 µg/ml for C2C12 cells. Infected cells were first propagated, then sorted by fluorescence-activated cell sorting (FACS) based on GFP expression. All cell lines were verified by immunofluorescence (IF) microscopy and Western blotting (WB) using protein tag specific antibodies.

### APEX2 proximity labeling

For conventional APEX-based proximity labeling, cells were incubated with 0.5 mM APEX substrate, biotin or desthiobiotin tyramide (BT, Iris Biotech, Cat. No. LS-3500.1000; or DTBT, Iris Biotech, LS-1660.0250; respectively), for 30 min at 37 °C before addition of H_2_O_2_ (Sigma Aldrich, Cat. No. H1009) to a final concentration of 1 mM and incubation at RT for 3 min. For DAAO-dependent (“iAPEX”) labeling, APEX substrate was added to the cells together with 10 mM D-amino acid at 37 °C for 30 min, unless noted otherwise. Unlabeled samples were kept untreated. After substrate incubation, the medium was aspirated, and cells washed three times with quenching buffer (1× PBS containing 10 mM sodium ascorbate, 10 mM sodium azide, and 5 mM Trolox). Cells grown on glass coverslips for IF microscopy were fixed immediately. Samples intended for proteomic and WB analyses were prepared by lysing and scraping the cells off the growth surface in ice-cold lysis buffer (0.5% [vol/vol] Triton X-100, 0.1% [wt/vol] SDS, 10% [wt/vol] glycerol, 300 mM NaCl, 100 mM Tris/HCl, pH 7.5, and protease inhibitors) containing 10 mM sodium ascorbate, 10 mM sodium azide, and 5 mM Trolox. The collected lysate was briefly vortexed, incubated on ice for 15 min, and cleared by centrifugation (20.000 g for 30 min at 4 °C).

### O_2_ consumption analysis

For the assay, 15.000 IMCD3 cells were seeded into a 96-well XF cell culture microplate (Agilent, Cat. No. 103794-100) in 80 µl of growth medium and left to settle at RT for 1 h. Cells were grown overnight at 37°C in a 5% CO₂ incubator, followed by serum starvation for 24 h. The sensor cartridge (Agilent, 103793-100) was prepared following the manufacturer’s instructions.

On the day of the assay, Seahorse XF Assay Medium (pH 7.4, Agilent, Cat. No. 103575-100) was prepared by adding Seahorse XF glucose (f.c. 17.5 mM, Agilent, Cat. No. 103577-100), pyruvate (f.c. 1 mM, Agilent, Cat. No. 103578-100) and L-glutamine (f.c. 2 mM, Agilent, Cat No. 103579-100). The cells were washed twice with 100 µl of prewarmed XF Assay Medium. Finally, 180 µl of XF Assay Medium was added to each well. The plate was incubated at 37°C in a non-CO₂ incubator for 60 minutes before starting the assay. Oligomycin (15 µM stock, f.c. 1.5 µM, Sigma-Aldrich, Cat. No. 4876), D- or L-amino acids (100 mM stock, f.c. 10 mM), and Hoechst (100 µM stock, f.c. 10 µM) were loaded into individual injection ports of the sensor cartridge. If a chemical was to be omitted during injection, medium was added to the designated port instead. The microplate and sensor cartridge were loaded into the Agilent Seahorse Analyzer. The experimental protocol included an initial calibration and equilibration step, and measurement cycles consisting of 3 min mixing, 15 s wait period and 3 min of measurement. O_2_ consumption rates (OCR) were measured for 20 cycles. Oligomycin and amino acids were injected after 4 and 8 cycles respectively. After 20 cycles Hoechst staining was performed for 3 min.

Wells with initial OCR values (Y1 rate) > 20 pmol/min, initial O₂ levels (Y1 level) ≈ 100 mmHg reducing to ≤ 20 mmHg after oligomycin, and initial pH near 7.4 were deemed acceptable, while those failing to meet these criteria were flagged and excluded during data analysis using the software’s plate map modification tool. Post-assay, the XF cell culture microplate was transferred to the BioTek Cytation system to allow normalization of each well to the respective cell number.

### Streptavidin affinity chromatography

Desthiobiotin tyramide labeled lysates were prepared as input for chromatography by adjusting them to equal concentrations and volumes. Fractions of input samples were taken as SDS-PAGE and WB controls. Streptavidin Sepharose High Performance Medium (Cytiva, 17-5113-01) was washed, equilibrated and then incubated with lysates at RT under rotation for 1 h. Unbound material was collected from settled beads and kept for WB analysis. Loaded beads were washed extensively with lysis buffer and spun dry before elution. Competitive elution buffer (100 mM Tris/HCl pH 7.5, 5 mM biotin) was added to the beads and incubated for 30 min shaking in a thermomixer at 950 rpm at RT. Elution was repeated and the eluates were combined. Eluates were subjected to centrifugal filter unit (Amicon Ultra, 0.5 ml 30K, Sigma-Aldrich, Cat. No. UFC5030) to be concentrated before mass spectrometric sample preparation. 10% of each eluate was kept for WB analysis.

### Mass spectrometry

For mass spectrometry, 12 x 10^6^ cells were seeded per 500 cm^2^ plate, grown for three days and starved for 24 h before APEX labeling and streptavidin chromatography. Mass spectrometry was performed at the EMBL Proteomic Core Facility in Heidelberg, Germany. For the mass spectrometric analysis, Amicon-concentrated eluates were subjected to an in-solution tryptic digest, following a modified version of the Single-Pot Solid-Phase-enhanced Sample Preparation (SP3) technology^79,80^. 20 µl of a slurry of hydrophilic and hydrophobic Sera-Mag Beads (Thermo Scientific, #4515-2105-050250, 6515-2105-050250) were mixed, washed with water and were then reconstituted in 100 µl water. 5 µl of the prepared bead slurry were added to 50 µl of the eluate following the addition of 55 µl of acetonitrile. All further steps were prepared using the King Fisher Apex System (Thermo Scientific). After binding to beads, beads were washed three times with 100 µl of 80% ethanol before they were transferred to 100 µl of digestion buffer (50 mM HEPES/NaOH pH 8.4 supplemented with 5 mM TCEP, 20 mM chloroacetamide (Sigma-Aldrich, #C0267), and 0.25 µg trypsin (Promega, #V5111). Samples were digested over night at 37°C, beads were removed, and the remaining peptides were dried down and subsequently reconstituted in 10 µl of water. 80 µg of TMT10plex (Thermo Scientific, #90111)^81^ label reagent dissolved in 4 µl of acetonitrile were added and the mixture was incubated for 1 h at room temperature. Excess TMT reagent was quenched by the addition of 4 µl of an aqueous solution of 5% hydroxylamine (Sigma, 438227). Mixed peptides were subjected to a reverse phase clean-up step (OASIS HLB 96-well µElution Plate, Waters #186001828BA). Peptides were subjected to an offline fractionation under high pH conditions^80^. The resulting 12 fractions were analyzed by multistage mass spectrometry (MS^3^) on a Lumos system (Thermo Scentific).

To this end, peptides were separated using an Ultimate 3000 nano RSLC system (Dionex) equipped with a trapping cartridge (Precolumn C18 PepMap100, 5 mm, 300 μm i.d., 5 μm, 100 Å) and an analytical column (Acclaim PepMap 100. 75 × 50 cm C18, 3 mm, 100 Å) connected to a nanospray-Flex ion source. The peptides were loaded onto the trap column at 30 µl per min using solvent A (0.1% formic acid) and eluted using a gradient from 2 to 80% Solvent B (0.1% formic acid in acetonitrile) over 2 h at 0.3 µl per min (all solvents were of LC-MS grade). The Orbitrap Fusion Lumos was operated in positive ion mode with a spray voltage of 2.5 kV and capillary temperature of 275 °C. Full scan MS spectra with a mass range of 375–1.500 m/z were acquired in profile mode using a resolution of 120.000 (maximum fill time of 50 ms; AGC Target was set to Standard) and a RF lens setting of 30%. Ions were selected in the quadrupole applying an isolation window of 1.5 m/z. Fragmentation was triggered by HCD using a collision energy of 36%. Ions were analyzed in the ion trap (maximum fill time of 50 ms; AGC target was set to Standard). The top 5 precursors were selected by synchronous precursor selection (SPS) between 400 to 2.000 m/z with a precursor ion exclusion width of -18 and +5 m/z. Their fragmentation was triggered by HCD using a collision energy of 70%. Ions were analyzed in the Orbitrap using a resolution of 50.000 (maximum injection time was set to 105 ms and the AGC Target was set to Custom and 200%).

Acquired data were analyzed using FragPipe^82^ and a Uniprot *Mus musculus* FASTA database (UP000000589, ID10090 with 21.968 entries, date: 27.10.2022, downloaded: January 11th 2023) including common contaminants. The following modifications were considered: Carbamidomethyl (C, fixed), TMT10plex (K, fixed), Acetyl (N-term, variable), Oxidation (M, variable) and TMT10plex (N-term, variable). The mass error tolerance for full scan MS spectra was set to 10 ppm and for MS/MS spectra to 0.02 Da. A maximum of 2 missed cleavages were allowed. A minimum of 2 unique peptides with a peptide length of at least seven amino acids and a false discovery rate below 0.01 were required on the peptide and protein level^83^.

### MS data analysis

Obtained TMT reporter intensities in one experiment were preprocessed using Perseus (version 2.0.11.0). Raw TMT signal intensities were log2 transformed, filtered for valid values in at least two out of three replicates in each group and missing values were replaced from normal distribution (width: 0.3; standard deviation down shift: 1.8). The filtered and imputed data was then Z-score normalized, where the mean of each column (reporter intensities for all proteins) was subtracted from each value and the result divided by the standard deviation of the column. Hierarchical cluster analyses were performed on the preprocessed and normalized data according to Ward’s minimum variance method using two-way unstandardized clustering in JMP software (Statistical Analysis System; v17.2.0). Candidates with *p* values < 0.05 were not displayed in clusters.

Gene Ontology enrichment analysis was performed using a web-based tool, the EnrichR (https://maayanlab.cloud/Enrichr/). Protein lists to be investigated (such as proteins in subclusters) were compared to all proteins identified in the respective mass spectrometry experiment as background list^84^.

### Expansion microscopy

Ultrastructure expansion microscopy (U-ExM) was performed following a modified protocol by Gambarotto *et al.*^85^ to achieve high-resolution imaging of physically magnified cellular structures. 5 × 10^4^ cells were seeded per well on round 12-mm #1.5 coverslips (Fisher Scientific, 11846933) in 24-well plates. APEX proximity labeling was performed as described. After quenching, 300 µl cross-linking solution (1.4% formaldehyde (Sigma-Aldrich, F8775) / 2% acrylamide (Sigma-Aldrich, A4058) in 1x PBS) was added to each well, and cells were incubated for 5 h at 37 °C. For gelation, 90 µl monomer solution mixed with 5 µl 10% APS (Thermo Fisher, 17874) and 5 µl 10% TEMED (Thermo Fisher, 17919) was drop-wise applied on parafilm placed in an ice-cold humid chamber. For 1 ml of monomer solution 500 µl of sodium acrylate (Sigma-Aldrich, 408220, 38% stock in nuclease-free water, f.c. 19%), 250 µl acrylamide (Sigma-Aldrich, A4058, 40% stock, f.c. 10%), 50 µl N,N’-methylenbisacrylamide (Sigma-Aldrich, M1533, 2% stock, fc 0.1%) and 100 µl 10x PBS were mixed. Coverslips were positioned cell-side down on the gelation solution, incubated on ice for 5 min, then at 37°C for 1 h. Gels were detached by transferring coverslips to 6-well plates with 1 ml denaturation buffer (200 mM SDS, 200 mM NaCl, 50 mM Tris/HCl in water, pH 9) for 15 min at RT, then incubated in 1.5 ml reaction tubes with fresh denaturation buffer at 95 °C for 1.5 h. For expansion, gels were transferred to individual 250 ml beakers with dH₂O and agitated at RT. Every 10 min water was replaced for three times. After measuring expansion, gels were shrunk in 1x PBS for 30 min and cut into quarters. For antibody staining, gel pieces were incubated in 200 µl of primary antibody diluted in 2% BSA (Carl Roth, 3737.3) in 1x PBS at 4 °C overnight. The next day, gels were washed with PBS + 0.1% Tween-20 (Carl Roth, 9127.2), then incubated in 200 µl of secondary antibody in 2% BSA in 1x PBS for 2.5 h at 37 °C with agitation, protected from light. After three washes gels were re-expanded in dH₂O with 0.02% sodium azide and incubated at 4 °C for three days.

25 mm coverslips were coated with poly-D-lysine (100 µg/ml (Gibco, A3890401) overnight at 4 °C, rinsed, dried, and stored. Imaging was conducted on a Zeiss LSM900 confocal microscope with AiryScan using a 63× oil objective (Plan-Apochromat 63x/1.40 Oil DIC M27), with the convex gel side placed against the coated coverslip surface in a round imaging chamber.

### Immunofluorescence microscopy

For fluorescence microscopy, all steps were performed at RT. 5 x 10^4^ cells were grown on round 12-mm #1.5 coverslips (Fisher Scientific, 11846933) in a 24-well plate and fixed in 4% PFA for 15 min. Cells were rinsed three times with 1x PBS between subsequent steps. Cells were incubated in blocking buffer (3% BSA, 5% serum, in 1x PBS) for 30 min. Primary and secondary antibody dilutions were prepared in blocking buffer. Primary antibody mixes were incubated on cells for 1 h, secondary antibody mixes for 30 min. The latter contained fluorescent secondary antibodies or streptavidin. DNA was stained with Hoechst 33258 (1:1000 in 1x PBS, Thermo Scientific, H3570). Lastly, coverslips were mounted on glass slides using Fluoromount G mounting medium (Fisher Scientific, 15586276). APEX2 enzyme was detected by GFP fluorescence, DAAO and fusion enzymes were detected by FLAG or ALFA staining.

Prepared specimens were imaged on Leica DMi8 microscope (LAS X software, version 3.7.0.20979) with PlanApochromat oil objectives (63×, 1.4 NA) using appropriate filters. Images were captured using a Leica DFC3000 G camera system. Images were processed using ImageJ2 (v2.14.0/1.54f).

### Primary cilia intensity quantification

Fluorescence intensities were quantified by ImageJ plugin CiliaQ (v0.1.4)^86^. A cilia mask was obtained by applying the RenyiEntropy threshold algorithm to either the ARL13B or GFP channel of each image.

### Live-Cell Imaging of Peroxidase-Catalyzed Amplex™ UltraRed Oxidation

Cells were seeded onto round 25 mm #1.5 coverslips (Fisher Scientific, 10593054) in a 6-well plate and serum-starved one day before imaging. Cells were imaged in live using either a Zeiss LSM800 with a 40x objective (Plan-Apochromat 40x/1.3 Oil DIC UV-IR M27) or a Zeiss LSM900 confocal microscope with a 63× oil objective (Plan-Apochromat 63x/1.40 Oil DIC M27), diode lasers 488 and 561 and appropriate filters. For imaging, the coverslip was placed in a round chamber and covered with cold (4 °C) HEPES-buffered DMEM/F12 medium without phenol red.

For live-cell time series of resorufin channel only, multi-channel images were taken before and after the Amplex UltraRed (AmUR) oxidation experiment. Resorufin channel recording was started before substrate addition to the cells and set to record as fast as possible. Cells were covered with 800 µl imaging buffer. AmUR (Fisher Scientific, 10737474) was pre-mixed with either D-Met or H_2_O_2_ to be added to the solution dome to a final concentration of 50 µM AmUR and 10 mM D-Met or 10 mM H_2_O_2_ respectively. Substrates and mixes were kept on ice.

For time-lapse imaging, 400 µl imaging medium was supplemented with 100 µM AmUR and 20 mM D-Met or 2 mM H_2_O_2_, then directly and carefully pipetted into the center of the 400 µl solution dome to yield halved final concentrations. Both GFP and resorufin channels were monitored at 4.8-second intervals over a total duration of 264 s.

### SDS-PAGE and Western blotting

For SDS-PAGE and Western blotting, standard techniques were applied. Cell lysates were generated as described before. 20 µg protein was separated on 4-12% Bis-Tris polyacrylamide gels (Invitrogen NuPAGE, WG1403BX10) and transferred onto nitrocellulose membrane (Fisher Scientific, 15269794) for fluorescence detection. Before blocking, membranes were dried and total protein stain (LI-COR. 926-11011) was used according to manufacturer’s guidelines to verify equal gel loading and transfer. Destained or untreated membranes were blocked in Intercept (TBS) Protein-free blocking buffer (LI-COR, 927-80001) at RT for 30 min. Specific primary antibody mixes were prepared in 5% milk in 1x TBS and membranes were incubated at 4 °C overnight. Fluorescently coupled secondary antibodies were used to visualize stained proteins and were imaged on a LI-COR Odyssey CLx laser scanner.

### Xenopus methods

All animals were treated according to the German regulations and laws for care and handling of research animals, and experimental manipulations were approved by the Regional Government Stuttgart, Germany (RPS35-9185.81/0471 and RPS35-9185-99/426).

For expression of iAPEX enzymes, capped mRNA was synthesized from linearized plasmids using mMessage mMachine kit (Invitrogen AM1344). Injection drop size was calibrated to 4 nl to deliver 300 ng of mRNA per single injection. Localization of constructs was analyzed in embryos fixed in 4 % PFA in 1× PBS for 1 h at room temperature or overnight at 4°C, followed dissection as indicated and IF as previously published^87^. For *in vivo* proximity labeling of ependymal cilia at st. 45, labeling solution and H_2_O_2_ were sequentially injected into the ventricular system. For post-fixation labeling, embryos were PFA-fixed for 20 min and dissected brains were incubated in BT and D-Norv for up to 10 min.

### Miscellaneous

All graphs were prepared with Graphpad Prism (10.3.1(464)). Figures were prepared using Affinity Designer (1.10.8).

### Antibodies and reagents

The following antibodies with indicated dilutions were used in this study:

| Antibody (clone) | Supplier/ CatN | Species | IgG | Clonality | Dilution<br>IF | Dilution<br>WB |
| --- | --- | --- | --- | --- | --- | --- |
| acTub (6-11B-1) | Sigma-Aldrich<br>(Merck) T7451 | mouse | IgG2b | monoclonal | 1:2000 | 1:2000 |
| ALFA sdAb<br>rabbit-fc | NanoTag<br>Biotechnologies<br>N1583 | camelid | IgG | polyclonal | 1:500 | 1:500 |
| ALFA sdAb mouse-fc | NanoTag Biotechnologies N1582 | camelid | IgG1 | monoclonal | 1:500 | 1:500 |
| Annexin V | Proteintech 66245-1-Ig | mouse | IgG1 | monoclonal | n.a. | 1:10000 |
| ARL13B | Proteintech 17711-1-AP | rabbit | IgG | polyclonal | 1:1000 | n.a. |
| FLAG (M2) | Sigma-Aldrich (Merck) F1804 | mouse | IgG1 | monoclonal | 1:1000 | 1:1000 |
| GAPDH (1E6D9) | Proteintech 60004-1-Ig | mouse | IgG2b | monoclonal | n.a. | 1:2000 |
| GFP (3H9) | Proteintech (Chromotek) 3h9 | rat | IgG2a | monoclonal | n.a. | 1:1000 |
| "GFP34" | self-made | rabbit | IgG | polyclonal | n.a. | 1:500 |
| IFT20 | Proteintech 13615-1-AP | rabbit | IgG | polyclonal | 1:2500 | n.a. |
| IFT57 | Proteintech 11083-1-AP | rabbit | IgG | polyclonal | n.a. | 1:500 |
| IFT88 | Proteintech 13967-1-AP | rabbit | IgG | polyclonal | 1:200 | 1:1000 |

| **Antibody** | **Supplier/ CatN** | **Host** | **Reac-tivity** | **Target Isotype** | **Dilution IF** | **Dilution WB** |
| --- | --- | --- | --- | --- | --- | --- |
| FluoTag®-X2 α-ALFA Alexa Fluor™ 647 | NanoTag Biotechnologies N1502-AF647-L | camelid sdAb | ALFA | n.a. | 1:500 | n.a. |
| FluoTag®-X2 α-ALFA IRdye® 800CW | NanoTag Biotechnologies N1502-Li800-L | camelid sdAb | ALFA | n.a. | n.a. | 1:1000 |
| Streptavidin Alexa Fluor™ 647 | Fisher Scientific S21374 | n.a. | biotin | n.a. | 1:1000 | n.a. |
| Streptavidin IRDye® 680RD | LI-COR 926-68079 | n.a. | biotin | n.a. | n.a. | 1:1000 |
| Streptavidin IRDye® 800CW | LI-COR 926-32230 | n.a. | biotin | n.a. | n.a. | 1:1000 |
| Streptavidin Northern-Lights™ 557 | RD Systems NL999 | n.a. | biotin | n.a. | n.a. | 1:2000 |
| α-mouse IgG HRP conjugate | Merck AP127P | goat | mouse | IgG | n.a. | 1:2000 |
| α-mouse IgG1 Alexa Fluor™ 488 | Thermo Fisher Scientific A-21121 | goat | mouse | IgG1 | 1:500 | n.a. |
| α-mouse IgG1 Cyanine3 | Thermo Fisher Scientific M30010 | goat | mouse | IgG1 | 1:500 | n.a. |
| α-mouse IgG1 Cyanine5 | Thermo Fisher Scientific A10524 | donkey | mouse | IgG | 1:500 | n.a. |
| α-mouse IgG2a Alexa Fluor™ 350 | Thermo Fisher Scientific A-21130 | goat | mouse | IgG2a | 1:500 | n.a. |
| α-mouse IgG2a<br>Alexa Fluor™ 488 | Thermo Fisher<br>Scientific A-21131 | goat | mouse | IgG2a | 1:500 | n.a. |
| α-mouse IgG2a<br>Alexa Fluor™ 555 | Thermo Fisher<br>Scientific A-21137 | goat | mouse | IgG2a | 1:500 | n.a. |
| α-mouse IgG2b<br>Alexa Fluor™ 488 | Thermo Fisher<br>Scientific A-21141 | goat | mouse | IgG2b | 1:500 | n.a. |
| α-mouse IRDye®<br>680RD | LI-COR 926-68070 | donkey | mouse | IgG | 1:500 | n.a. |
| α-rabbit IgG Alexa<br>Fluor™ 350 | Thermo Fisher<br>Scientific A-11046 | goat | rabbit | IgG | 1:500 | n.a. |
| α-rabbit IgG Alexa<br>Fluor™ 488 | Thermo Fisher<br>Scientific A-11008 | donkey | rabbit | IgG | 1:500 | n.a. |
| α-rabbit IgG Alexa<br>Fluor™ 555 | Thermo Fisher<br>Scientific A-21428 | donkey | rabbit | IgG | 1:500 | n.a. |
| α-rabbit IgG Alexa<br>Fluor™ 647 | Thermo Fisher<br>Scientific A-21245 | donkey | rabbit | IgG | 1:500 | n.a. |
| α-rabbit IgG Alexa<br>Fluor™ plus555 | Thermo Fisher<br>Scientific A-31572 | donkey | rabbit | IgG | 1:500 | n.a. |
| α-rabbit IgG HRP-conjugate | Sigma-Aldrich<br>AP187P | goat | rabbit | IgG | n.a. | 1:2000 |
| α-rabbit IRDye®<br>800CW | LI-COR 926-32211 | donkey | rabbit | IgG | n.a. | 1:10000 |
| α-rat Cyanine5 | Thermo Fisher<br>Scientific<br>A10525 | donkey | rat | IgG | 1:500 | n.a. |
| α-rat IgG IRDye®<br>680RD | LI-CO 926-68076 | goat | rat | IgG | n.a. | 1:10000 |
| α-rat IgG IRDye®<br>800CW | LI-COR 926-32219 | goat | rat | IgG | n.a. | 1:10000 |

### Cell lines used in this study

| Cell line | Parental cell line | reference |
| --- | --- | --- |
| cilia-APEX2 | IMCD3 FlpIn | May et al., 2021<sup>17</sup> |
| cyto-APEX2 | IMCD3 FlpIn | May et al., 2021<sup>17</sup> |
| *Cep164*<sup>-/-</sup> cilia-APEX2 | IMCD3 FlpIn | May et al., 2021<sup>17</sup> |
| cilia-iAPEX | IMCD3 FlpIn | This work |
| cyto-iAPEX<br>(cilia-APEX2, cyto-DAAO) | IMCD3 FlpIn | This work |
| cilia-DAAO-APEX2 | IMCD3 FlpIn | This work |
| cilia-iAPEX | NIH/3T3 FlpIn | This work |
| cilia-iAPEX | RPE FRT/TO | This work |
| cilia-iAPEX (virus) | IMCD3 FlpIn | This work |
| cilia-iAPEX | 3T3-L1 | This work |
| cilia-iAPEX | C2C12 | This work |
| NIH/3T3 | - | gift from P. Niewiadomski |
| 3T3-L1 | - | gift from D. Wachten |
| C2C12 | - | ATCC |
| RPE1 FRT/TO | - | gift from J. Mansfeld |
| HEK293T | - |  |

## ACKNOWLEDGEMENTS

We thank D. Yildiz for fluorescence-activated cell sorting, P. Niewiadomski, D. Wachten, J. Mansfeld, and L. Prates Roma for reagents, D. Jann and S. Plant for experimental assistance, D.K. Breslow and B. Schrul for helpful discussions and comments on the manuscript. We thank all members of the Mick lab for stimulating discussions. This work was supported by Deutsche Forschungsgemeinschaft (DFG) funding to D.U.M. (TRR152-P28 – Project-ID 239283807, FOR5547-P3 – Project-ID 503306912, and Project-ID 513767027). We are grateful to F. Stein at the EMBL proteomics core facility for MS data analysis.

## AUTHOR CONTRIBUTIONS

Conceptualization, TJS and DUM; Methodology, TJS, LKS, AP, PH, KvdM, VC, JH and KF; Formal analysis, TJS, LKS, AP, KvdM, VC, and KF; Writing – original draft, TJS and DUM; Writing – review & editing, TJS, KF, KvdM, AP and DUM; Visualization, TJS, LKS, KF, AP and VC; Funding acquisition, DUM; Supervision, KF, and DUM.

---

**Fig. S1:**
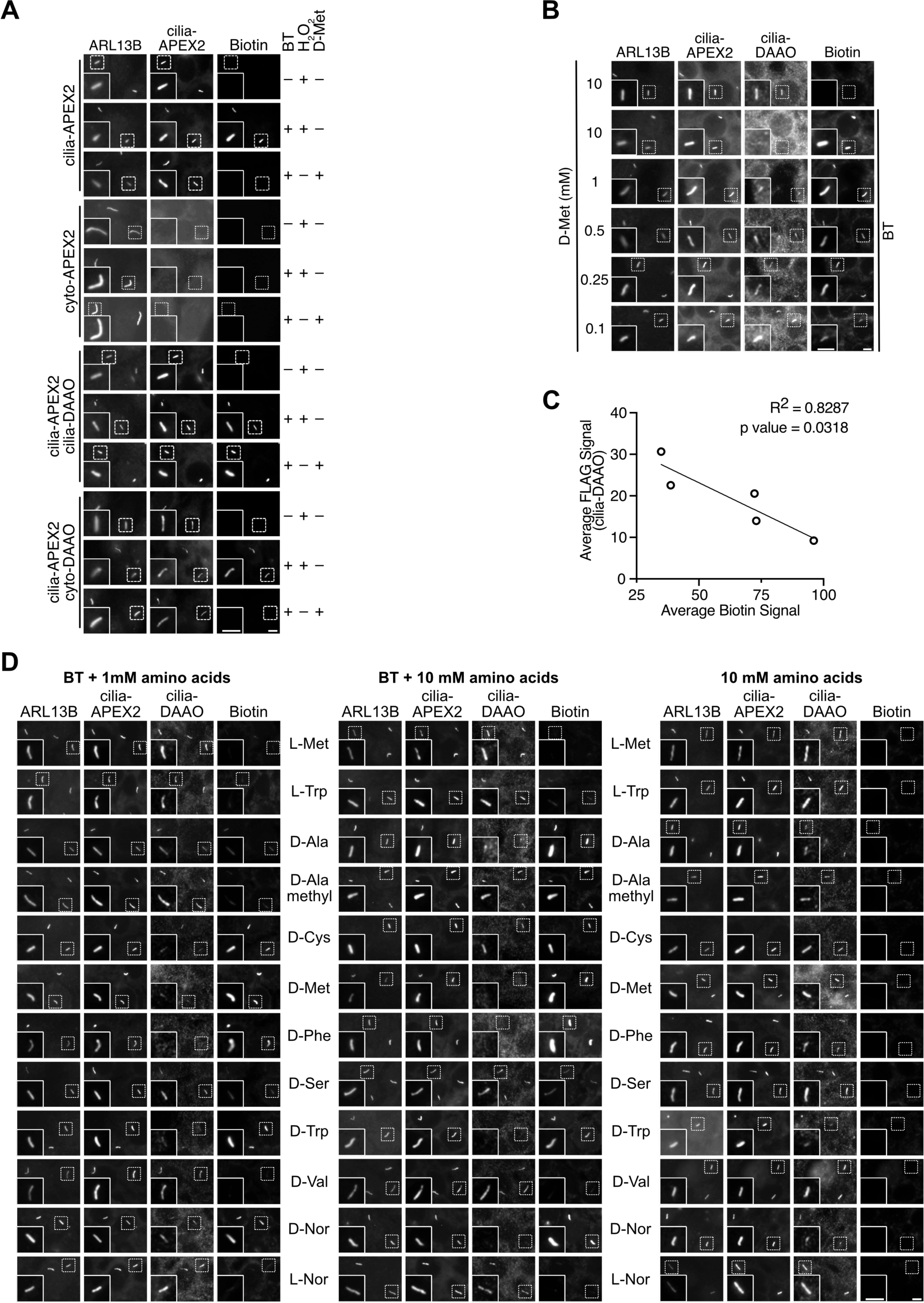
The DAAO-APEX2 enzymatic cascade requires co-localization and can use various D-amino acids as substrates. **(A)** Representative IF images of IMCD3 cells expressing cilia-APEX2, cilia-iAPEX and the cytosolic controls, cyto-APEX and cyto-DAAO. Cells were incubated with biotin tyramide (BT) for 30 min together with H_2_O_2_ for 3 min for direct activation of APEX2, or with D-Met for 30 min for local production of H_2_O_2_ by DAAO as indicated. ARL13B antibody was used to stain for primary cilia. GFP fluorescence marked APEX2. Biotin was detected by fluorescent streptavidin. **(B)** Proximity biotinylation of FLAG epitope precludes antibody binding. Proximity labeling in cilia-iAPEX IMCD3 cells was performed with indicated D-Met concentrations and cells analyzed by immunofluorescence microscopy, using fluorescently labeled streptavidin to detect biotin, anti-FLAG antibodies to detect cilia-DAAO, and anti-ARL13B antibodies to detect cilia. GFP fluorescence visualizes cilia-APEX2. **(C)** FLAG and biotin signals from micrographs as shown in (B) have been quantified and averages plotted. **(D)** Representative IF images of D-amino acid-dependent proximity labeling in IMCD3 cells expressing cilia-iAPEX. Various D- and L-amino acids were incubated at 1 or 10 mM concentration together with or without 500 µM BT for 30 min. ARL13B antibody was used to stain for primary cilia. GFP fluorescence marked APEX2. FLAG antibody marked cilia-DAAO. Biotin was detected by fluorescent streptavidin. All scale bars = 5 μm.

**Fig. S2:**
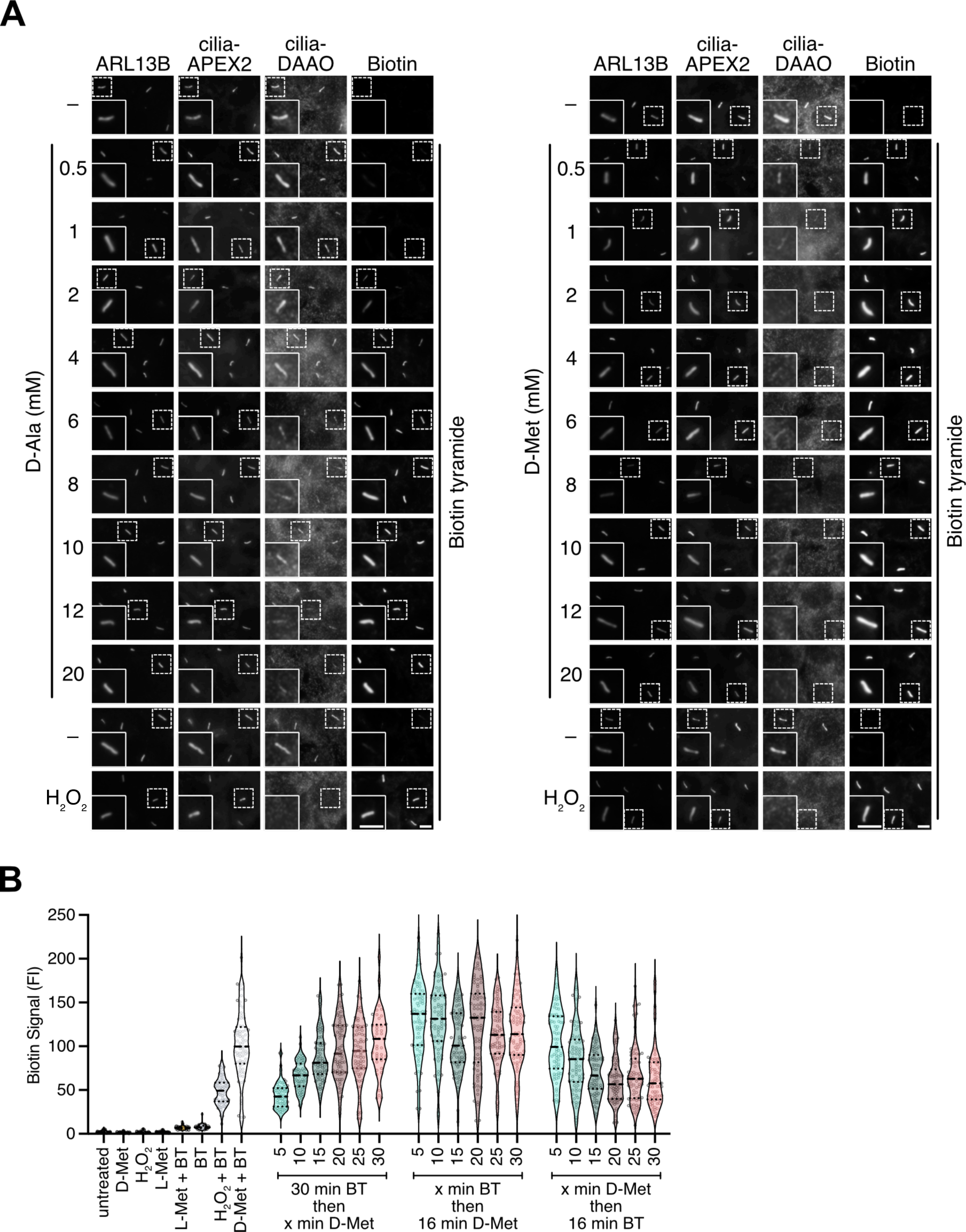
DAAO in vicinity of APEX2 allows spatiotemporal proximity labeling with amino acid substrate and concentration dependence. **(A)** The iAPEX-based biotinylation relies on DAAO-mediated oxidative deamination and is substrate and concentration dependent. Representative IF micrographs of DAAO substrate titration in IMCD3 cells stably expressing cilia-iAPEX (quantification shown in **Fig. 2C**). APEX labeling was performed by incubating cells for 30 min with 500 μM biotin tyramide (BT) together with varying concentrations (range from 0.5-20 mM) of D-alanine or D-methionine or 1 mM H_2_O_2_ (for 2 min). Antibody staining against ARL13B labeled primary cilia, while staining against FLAG marked cilia-DAAO. GFP fluorescence visualized cilia-APEX2, and biotin was detected by fluorescent streptavidin. **(B)** Pre-incubation with BT increases (left) while pre-incubation with D-Met lowers APEX2 activity (right). Violin plots show the CiliaQ quantified ciliary biotin signals. APEX labelings were performed by incubating cells for different times and in different orders with 500 µM BT and 10 mM D- or L-Met. Quartiles and median are indicated by dotted and dashed lines, respectively. n = 77 cilia per condition. All scale bars = 5 µm.

**Fig. S3:**
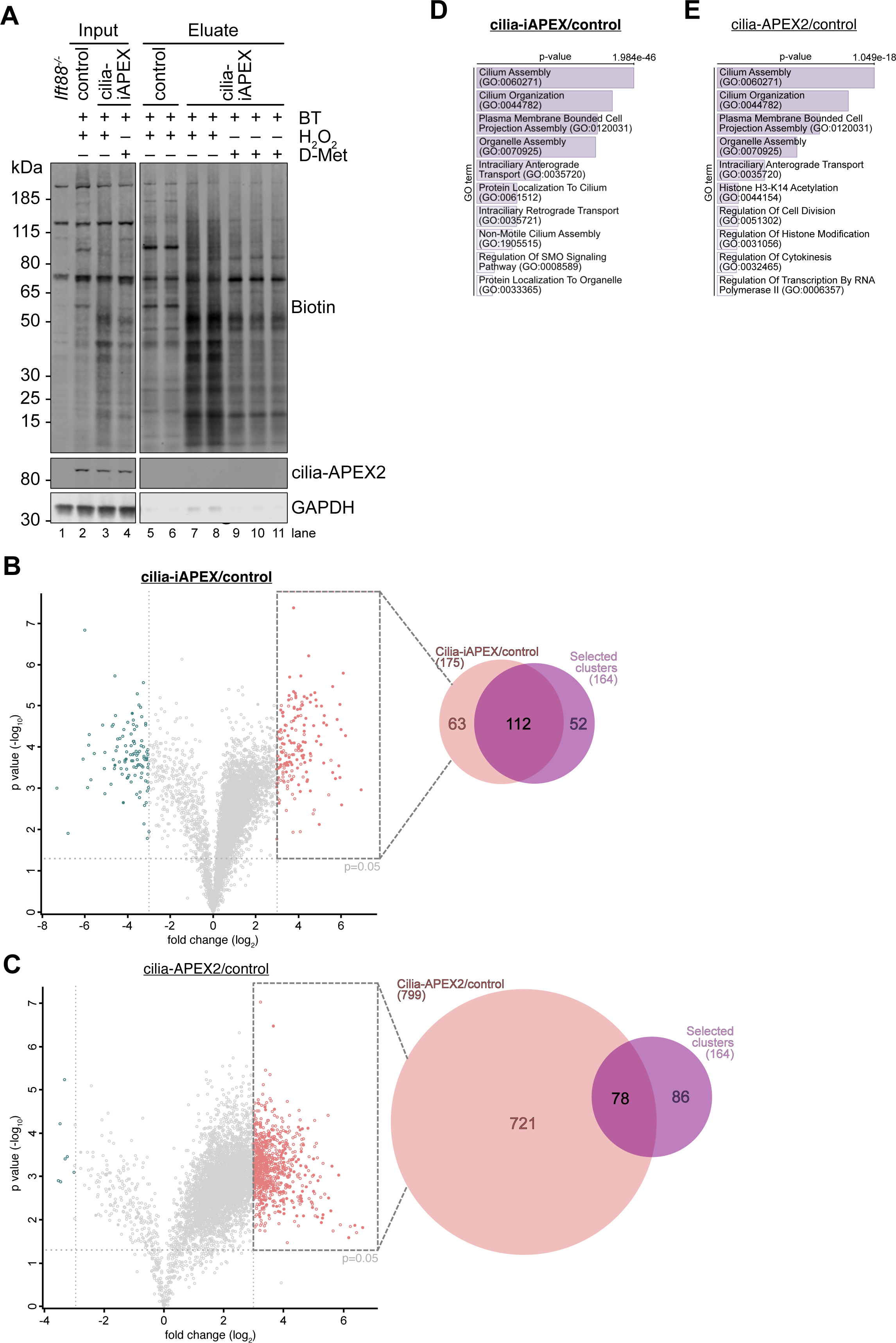
Hierarchical clustering of cilia-iAPEX proteomics data shows higher sensitivity and specificity than enrichment analysis. **(A)** Western blot analysis of samples after proximity labeling from IMCD3 cilia-iAPEX or cilia-ablated cilia-APEX2 *Cep164^-/-^* cells (control), as outlined in (**Fig. 4A**). Lysate from IMCD3 *Ift88^-/-^* cells served as an antibody specificity and untreated control (see **Fig. 4B**). Input and Eluate samples were separated by SDS-PAGE and analyzed by Western Blotting. Biotin was detected by fluorescently labeled streptavidin, cilia-APEX2 by antibodies against GFP. Input 0.063 %, Eluate 1.5 %. Stronger biotinylation was observed after H_2_O_2_-induced labeling. **(B and C)** Volcano plots display statistical significance *versus* protein enrichment of cilia-iAPEX2 **(B)** and cilia-APEX2 **(C)** proteomics compared with control samples. *p* values (unpaired Student’s *t* test) and TMT ratios were calculated from duplicate samples and plotted for 5982 proteins. Proteins are indicated by grey circles. Proteins with TMT ratios >2^3^ and <2^3^ are indicated by red and blue circles, respectively. Filled circles mark known cilia proteins identified in May et al., 2021^17^. Venn diagrams show numbers of enriched proteins (TMT ratios >2^3^) and overlap with selected cilia protein clusters from IMCD3 cells (see **Fig. 4D**) **(D)** GO_term enrichment analysis of cilia-iAPEX candidate proteins from (B) shows highly significant enrichment of proteins associated with cilia processes, including SHH signaling and protein trafficking. **(E)** GO_term enrichment analysis of proteins enriched in cilia-APEX2 samples (C) shows lower *p* values and enrichment of non-ciliary categories. *p* values were calculated by Fisher’s exact test.

**Fig. S4:**
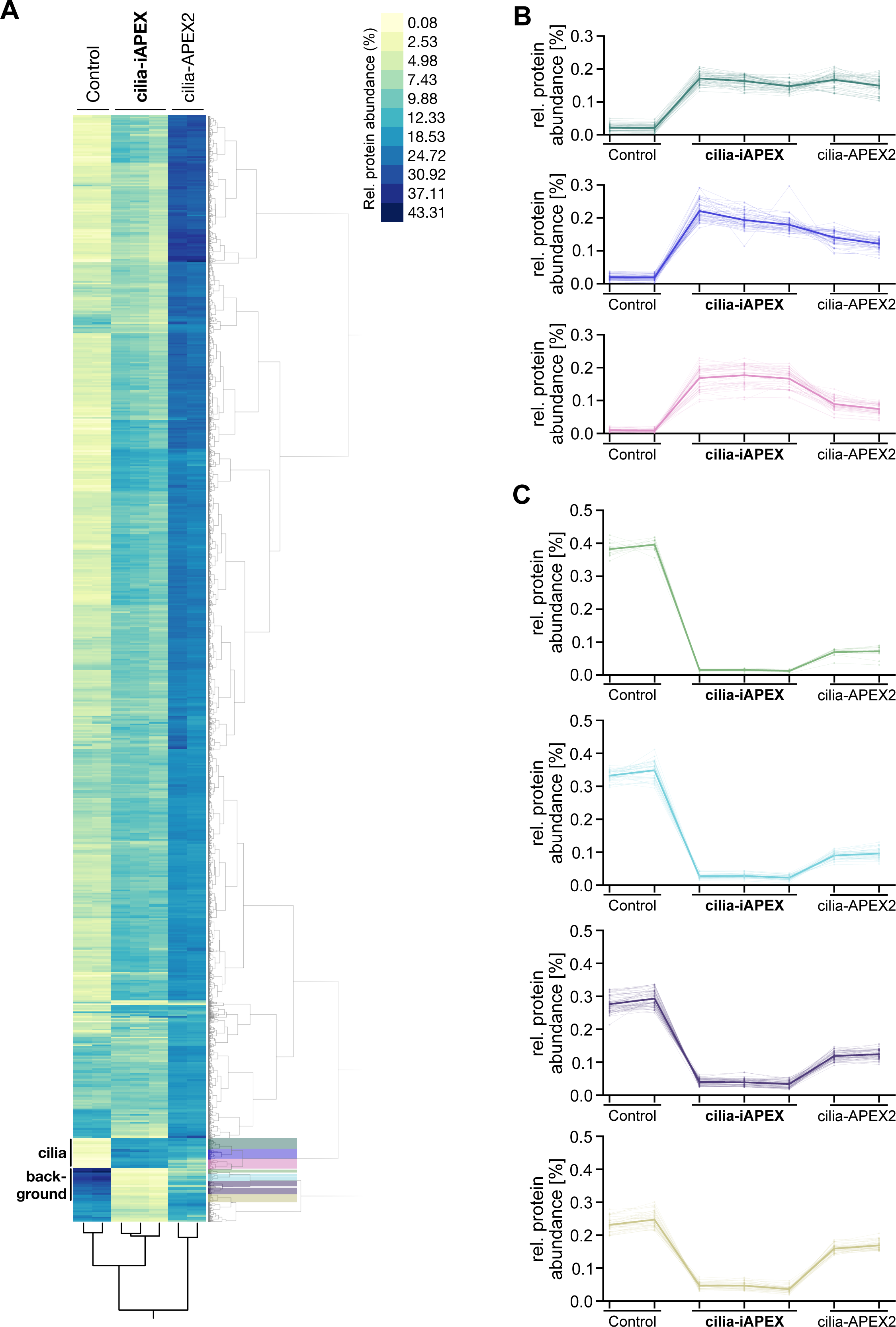
Hierarchical clustering of cilia-iAPEX proteomics in IMCD3 cells identifies cilia proteins as well as false-positive hits of previous studies. **(A)** Relative protein abundances (rows) of the individual samples (columns) from IMCD3 cilia-iAPEX proteomics experiment (see Fig. 4A) was analyzed by two-way hierarchical clustering (Ward’s method). Relative abundance of each protein was determined by dividing its individual TMT signal by the sum of TMT signals across all samples. The color legend for relative abundances (in %) is displayed. All quantified proteins are shown. Clusters containing cilia proteins, as well as example background clusters were highlighted. **(B** and **C)** Cilia clusters **(B)** and selected background clusters **(C)** are shown in magnified views. The average abundances of all proteins within the clusters are represented by thick lines, individual proteins by thin lines.

**Fig. S5:**
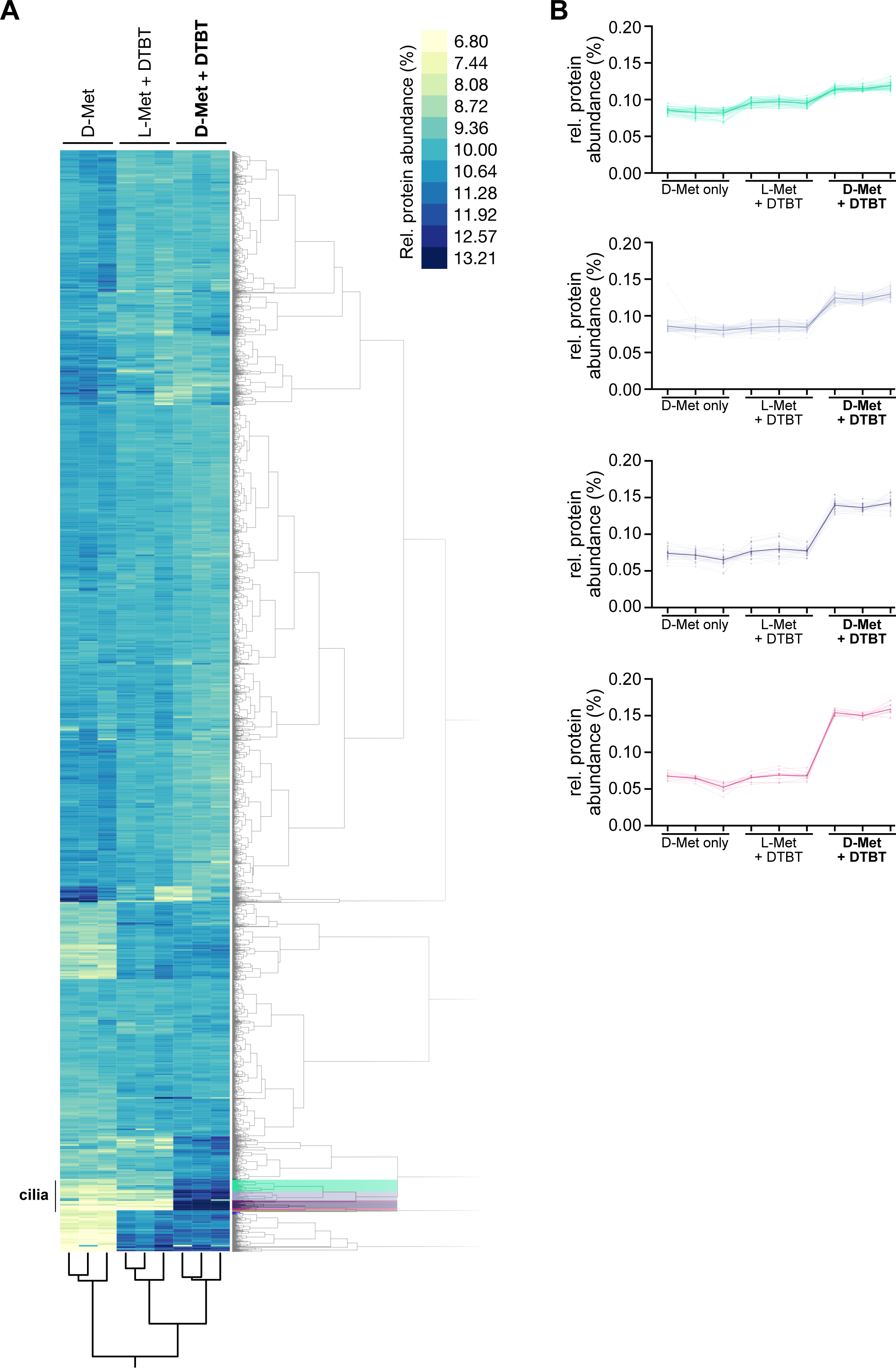
Hierarchical clustering of cilia-iAPEX proteomics in NIH/3T3 cells identifies cilia proteins. **(A)** Relative protein abundances (rows) of the individual samples (columns) from NIH/3T3 cilia-iAPEX proteomics experiment (see **Fig. 5A**) was analyzed by two-way hierarchical clustering (Ward’s method). Clusters containing cilia are indicated. All quantified proteins are shown. **(B)** Clusters containing cilia proteins are shown in magnified views, with thick lines showing average abundances of all proteins within the clusters and thin lines showing individual proteins.

## SUPPLEMENTARY TABLES

**Table S1: cilia-iAPEX proteomics of IMCD3 primary cilia**

First tab ‘Full IMCD3 Dataset’ lists the entire dataset from experiment, as depicted in Fig. 4A. Samples from mislocalized cyto-iAPEX cell lines were included in 10plex TMT experiment. UniProt identifiers, Gene names and protein descriptions according to the *Mus musculus* proteome database (UP000000589, ID10090, date: 27.10.2022) are listed. Column D shows number of unique quantified peptides for each protein. RAW TMT reporter intensities were summed in Column O to calculate relative TMT protein abundances. Imputed data marked in orange. Columns Z and AA display log_2_-transformed average TMT ratios and *p* values of cilia-iAPEX over *Cep164^-/-^* control, respectively. Columns AB and AC display log_2_-transformed average TMT ratios and *p* values of cilia-APEX2 over control, respectively. *p* values were calculated by two-sided Student’s *t* test.

Second tab ‘Extracted Cilia Clusters” displays data for proteins in cilia clusters shown in Fig. 4D. Colors highlight individual clusters as in **Fig. S4**. Average column means were calculated and displayed in **Fig. S4B**.

Third tab ‘Extracted Background Clusters” displays data for proteins in background clusters, parts of which are shown in **Fig. 4F**. Average column means were calculated for **Fig. S4C**.

Fourth tab ‘Legend’ explains columns in other tabs.

**Table S2: cilia-iAPEX proteomics of NIH/3T3 primary cilia**

First tab ‘Full NIH 3T3 Dataset’ lists the entire dataset from experiment depicted in Fig. 5A. Samples from cilia-iAPEX NIH/3T3 cells treated with D-Met (no BT control), BT+L-Met (L-Met control) and BT+D-Met (iAPEX) were included in a 10plex TMT experiment. UniProt identifiers, Gene names and protein descriptions according to the *Mus musculus* proteome database (UP000000589, ID10090, date: 27.10.2022) are listed. Column D shows number of unique quantified peptides for each protein. TMT reporter intensities were Z score column normalized in Columns O to X and summed in Column Y to calculate relative TMT protein abundances. Imputed data was marked in orange. Columns AJ and AK display log_2_-transformed average TMT ratios and *p* values of BT+D-Met over D-Met samples, respectively. Columns AL and AM display log_2_-transformed average TMT ratios and *p* values of cilia-APEX2 over control, respectively. *p* values were calculated by two-sided Student’s *t* test.

Second tab ‘Extracted Cilia Clusters” displays data for proteins in cilia clusters shown in Fig. 5D. Colors highlight individual clusters as in **Fig. S5**. Average column means were calculated and displayed in **Fig. S5B**.

Third tab ‘Legend’ explains columns in other tabs.

**Table S3: Comparison of cilia-iAPEX proteomics to cilia-APEX2**

First tab ‘May *et al*., 2021’ lists Gene names of the cilia-APEX2 proteome as defined in May *et al*., 2021.

Second tab ‘IMCD3 cilia-iAPEX’ lists proteins identified in three cilia clusters from IMCD3 cilia-iAPEX proteomics (**Fig. 4D** and **Table S1**).

Third tab ‘NIH-3T3 cilia-iAPEX’ lists Gene names of proteins identified in four cilia clusters from cilia-iAPEX proteomics in NIH/3T3 cells (selected clusters shown **Fig. 5D** and **Table S2**).

Fourth tab ‘Venn diagram’ lists Gene names in the individual Venn diagram sets and intersections (**Fig. 5E**) as indicated.

## SUPPLEMENTARY VIDEOS

**Video 1: H_2_O_2_ addition results in burst of peroxidase activity throughout the cell**

Live-cell imaging of cilia-iAPEX expressing IMCD3 cells. 50 µM AmUR and 10 mM H_2_O_2_ were added when indicated. Time lapse shows resorufin autofluorescence. Stills show cilia-APEX2 (detected by GFP autofluorescence) and resorufin fluorescence before and after substrate addition. (related to **Fig. 3C**)

**Video 2: D-Met-mediated DAAO activation results in specific peroxidase activity in cilia**

Live-cell imaging of cilia-iAPEX expressing IMCD3 cells. 50 µM AmUR and 10 mM D-Met were added when indicated. Time lapse shows resorufin autofluorescence. Stills show cilia-APEX2 (detected by GFP autofluorescence) and resorufin fluorescence before and after substrate addition. (related to **Fig. 3D**)

## REFERENCES

1. Qin, W., Cho, K. F., Cavanagh, P. E. & Ting, A. Y. Deciphering molecular interactions by proximity labeling. Nature Methods 1–11 (2021) doi:10.1038/s41592-020-01010-5.

2. Kang, M.-G. & Rhee, H.-W. Molecular Spatiomics by Proximity Labeling. Acc. Chem. Res. (2022) doi:10.1021/acs.accounts.2c00061.

3. Fedoryshchak, R. O. et al. Discovery of lipid-mediated protein–protein interactions in living cells using metabolic labeling with photoactivatable clickable probes. Chem. Sci. 14, 2419–2430 (2023).

4. Becker, A. P. et al. Lipid- and protein-directed photosensitizer proximity labeling captures the cholesterol interactome. bioRxiv 2024.08.20.608660 (2024) doi:10.1101/2024.08.20.608660.

5. Kalocsay, M. APEX Peroxidase-Catalyzed Proximity Labeling and Multiplexed Quantitative Proteomics. Methods Mol Biol 2008, 41–55 (2019).

6. Liu, X., Salokas, K., Weldatsadik, R. G., Gawriyski, L. & Varjosalo, M. Combined proximity labeling and affinity purification−mass spectrometry workflow for mapping and visualizing protein interaction networks. Nature Protocols 1–30 (2020) doi:10.1038/s41596-020-0365-x.

7. Mill, P., Christensen, S. T. & Pedersen, L. B. Primary cilia as dynamic and diverse signalling hubs in development and disease. Nat Rev Genet 1–21 (2023) doi:10.1038/s41576-023-00587-9.

8. Gopalakrishnan, J. et al. Emerging principles of primary cilia dynamics in controlling tissue organization and function. EMBO J 42, e113891 (2023).

9. Hilgendorf, K. I., Myers, B. R. & Reiter, J. F. Emerging mechanistic understanding of cilia function in cellular signalling. Nat Rev Mol Cell Biol 1–19 (2024) doi:10.1038/s41580-023-00698-5.

10. Moran, A. L., Louzao-Martinez, L., Norris, D. P., Peters, D. J. M. & Blacque, O. E. Transport and barrier mechanisms that regulate ciliary compartmentalization and ciliopathies. Nat Rev Nephrol 1–18 (2023) doi:10.1038/s41581-023-00773-2.

11. Su, S. et al. Genetically encoded calcium indicator illuminates calcium dynamics in primary cilia. Nat Methods 10, 1105–1107 (2013).

12. Ishikawa, H. & Marshall, W. F. Isolation of mammalian primary cilia. Methods Enzymol 525, 311–325 (2013).

13. Mick, D. U. et al. Proteomics of Primary Cilia by Proximity Labeling. Developmental Cell 35, 497–512 (2015).

14. Kohli, P. et al. The ciliary membrane-associated proteome reveals actin-binding proteins as key components of cilia. EMBO reports 18, e201643846–e201643846 (2017).

15. Arslanhan, M. D., Gulensoy, D. & Firat-Karalar, E. N. A Proximity Mapping Journey into the Biology of the Mammalian Centrosome/Cilium Complex. Cells 9, 1390 (2020).

16. Chen, X., Shi, Z., Yang, F., Zhou, T. & Xie, S. Deciphering cilia and ciliopathies using proteomic approaches. The FEBS Journal **n/a**, (2022).

17. May, E. A. et al. Time-resolved proteomics profiling of the ciliary Hedgehog response. Journal of Cell Biology 220, (2021).

18. Liu, X. et al. Numb positively regulates Hedgehog signaling at the ciliary pocket. Nat Commun 15, 3365 (2024).

19. Milione, R. R., Schell, B.-B., Douglas, C. J. & Seath, C. P. Creative approaches using proximity labeling to gain new biological insights. Trends in Biochemical Sciences (2023) doi:10.1016/j.tibs.2023.12.005.

20. Branon, T. C. et al. Efficient proximity labeling in living cells and organisms with TurboID. Nat Biotechnol 36, 880–887 (2018).

21. Lee, S.-Y. et al. Engineered allostery in light-regulated LOV-Turbo enables precise spatiotemporal control of proximity labeling in living cells. Nat Methods 20, 908–917 (2023).

22. Rhee, H.-W. et al. Proteomic Mapping of Mitochondria in Living Cells via Spatially-Restricted Enzymatic Tagging. Science 339, 1328–1331 (2013).

23. Eid, M., Barayeu, U., Sulková, K., Aranda-Vallejo, C. & Dick, T. P. Using the heme peroxidase APEX2 to probe intracellular H2O2 flux and diffusion. Nat Commun 15, 1239 (2024).

24. Mick, D. U. Establishing Cell Culture-Based Experimental Setups for Proximity Labeling Using Ascorbate Peroxidase (APEX). in Proximity Labeling (eds. Sunbul, M. & Jäschke, A.) vol. 2008 29–39 (Springer New York, New York, NY, 2019).

25. Saputra, F., Kishida, M. & Hu, S.-Y. Oxidative stress induced by hydrogen peroxide disrupts zebrafish visual development by altering apoptosis, antioxidant and estrogen related genes. Sci Rep 14, 14454 (2024).

26. Ransy, C., Vaz, C., Lombès, A. & Bouillaud, F. Use of H2O2 to Cause Oxidative Stress, the Catalase Issue. Int J Mol Sci 21, 9149 (2020).

27. Kisty, E. A., Falco, J. A. & Weerapana, E. Redox proteomics combined with proximity labeling enables monitoring of localized cysteine oxidation in cells. Cell Chemical Biology 30, 321–336.e6 (2023).

28. Molla, G., Motteran, L., Piubelli, L., Pilone, M. S. & Pollegioni, L. Regulation of D-amino acid oxidase expression in the yeast Rhodotorula gracilis. Yeast 20, 1061–1069 (2003).

29. Pollegioni, L., Piubelli, L., Sacchi, S., Pilone, M. S. & Molla, G. Physiological functions of D-amino acid oxidases: from yeast to humans. Cell. Mol. Life Sci. 64, 1373–1394 (2007).

30. Pollegioni, L. & Molla, G. New biotech applications from evolved D-amino acid oxidases. Trends in Biotechnology 29, 276–283 (2011).

31. Alim, I., Haskew-Layton, R. E., Aleyasin, H., Guo, H. & Ratan, R. R. Spatial, Temporal, and Quantitative Manipulation of Intracellular Hydrogen Peroxide in Cultured Cells. Methods Enzymol 547, 251–273 (2014).

32. Delling, M., Decaen, P. G., Doerner, J. F., Febvay, S. & Clapham, D. E. Primary cilia are specialized calcium signalling organelles. Nature (2013) doi:10.1038/nature12833.

33. Hashimoto, A. et al. Free d-serine, d-aspartate and d-alanine in central nervous system and serum in mutant mice lacking d-amino acid oxidase. Neuroscience Letters 152, 33–36 (1993).

34. Van Horn, M. R., Strasser, A., Miraucourt, L. S., Pollegioni, L. & Ruthazer, E. S. The Gliotransmitter d-Serine Promotes Synapse Maturation and Axonal Stabilization In Vivo. J Neurosci 37, 6277–6288 (2017).

35. Seckler, J. M. & Lewis, S. J. Advances in D-Amino Acids in Neurological Research. International Journal of Molecular Sciences 21, 7325 (2020).

36. Gambarotto, D. et al. Imaging cellular ultrastructures using expansion microscopy (U-ExM). Nat Methods 16, 71–74 (2019).

37. Den Toom, W. T. F., et al. Oxygen-consumption based quantification of chemogenetic H2O2 production in live human cells. Free Radical Biology and Medicine 206, 134–142 (2023).

38. Matlashov, M. E., Belousov, V. V. & Enikolopov, G. How much H(2)O(2) is produced by recombinant D-amino acid oxidase in mammalian cells? Antioxid Redox Signal 20, 1039–1044 (2014).

39. Zhou, M., Diwu, Z., Panchuk-Voloshina, N. & Haugland, R. P. A Stable Nonfluorescent Derivative of Resorufin for the Fluorometric Determination of Trace Hydrogen Peroxide: Applications in Detecting the Activity of Phagocyte NADPH Oxidase and Other Oxidases. Analytical Biochemistry 253, 162–168 (1997).

40. Martell, J. D., Deerinck, T. J., Lam, S. S., Ellisman, M. H. & Ting, A. Y. Electron microscopy using the genetically encoded APEX2 tag in cultured mammalian cells. Nat Protoc 12, 1792– 1816 (2017).

41. Hung, V. et al. Spatially resolved proteomic mapping in living cells with the engineered peroxidase APEX2. Nat Protoc 11, 456–475 (2016).

42. Del Olmo, T. et al. APEX2-mediated RAB proximity labeling identifies a role for RAB21 in clathrin-independent cargo sorting. EMBO reports 20, e47192 (2019).

43. McAlister, G. C. et al. MultiNotch MS3 Enables Accurate, Sensitive, and Multiplexed Detection of Differential Expression across Cancer Cell Line Proteomes. Anal. Chem. 86, 7150–7158 (2014).

44. Pappireddi, N., Martin, L. & Wühr, M. A Review on Quantitative Multiplexed Proteomics. Chembiochem 20, 1210–1224 (2019).

45. Götzke, H. et al. The ALFA-tag is a highly versatile tool for nanobody-based bioscience applications. Nat Commun 10, 4403 (2019).

46. Gómez, A. E., Christman, A. K., Weghe, J. C. V. D., Finn, M. & Doherty, D. Systematic analysis of cilia characteristics and Hedgehog signaling in five immortal cell lines. PLOS ONE 17, e0266433 (2022).

47. Kim, S. et al. Nde1-mediated inhibition of ciliogenesis affects cell cycle re-entry. Nat Cell Biol 13, 351–360 (2011).

48. Lai, C. K. et al. Functional characterization of putative cilia genes by high-content analysis. MBoC 22, 1104–1119 (2011).

49. Filipová, A. et al. Ionizing radiation increases primary cilia incidence and induces multiciliation in C2C12 myoblasts. Cell Biol Int 39, 943–953 (2015).

50. Rahimi, A. M., Cai, M. & Hoyer-Fender, S. Heterogeneity of the NIH3T3 Fibroblast Cell Line. Cells 11, 2677 (2022).

51. Bosch, J. A., Chen, C.-L. & Perrimon, N. Proximity-dependent labeling methods for proteomic profiling in living cells: an update. Wiley Interdiscip Rev Dev Biol 10, e392 (2021).

52. Paek, J. et al. Multidimensional Tracking of GPCR Signaling via Peroxidase-Catalyzed Proximity Labeling. Cell 169, 338–349.e11 (2017).

53. Finnegan, M. et al. Mode of action of hydrogen peroxide and other oxidizing agents: differences between liquid and gas forms. Journal of Antimicrobial Chemotherapy 65, 2108–2115 (2010).

54. Park, W. H. The effects of exogenous H2O2 on cell death, reactive oxygen species and glutathione levels in calf pulmonary artery and human umbilical vein endothelial cells. International Journal of Molecular Medicine 31, 471–476 (2013).

55. Kano, R. et al. In vivo cytosolic H2O2 changes and Ca2+ homeostasis in mouse skeletal muscle. *American Journal of Physiology-Regulatory*, Integrative and Comparative Physiology 326, R43– R52 (2024).

56. Genchi, G. An overview on d-amino acids. Amino Acids 49, 1521–1533 (2017).

57. Steinhorn, B. et al. Chemogenetic generation of hydrogen peroxide in the heart induces severe cardiac dysfunction. Nat Commun 9, 4044 (2018).

58. Kalinichenko, A. L. et al. Chemogenetic emulation of intraneuronal oxidative stress affects synaptic plasticity. Redox Biol 60, 102604 (2023).

59. Sorrentino, A., Eroglu, E. & Michel, T. Chapter 7 - In vivo applications of chemogenetics in redox (patho)biology. in *Oxidative Stress* (ed. Sies, H.) 97–112 (Academic Press, 2020). doi:10.1016/B978-0-12-818606-0.00007-9.

60. Rosini, E., Pollegioni, L., Ghisla, S., Orru, R. & Molla, G. Optimization of d-amino acid oxidase for low substrate concentrations – towards a cancer enzyme therapy. The FEBS Journal 276, 4921–4932 (2009).

61. Erdogan, Y. C. et al. Complexities of the chemogenetic toolkit: differential mDAAO activation by D-amino substrates and subcellular targeting. Free Radic Biol Med 177, 132–142 (2021).

62. Zaragozá, R. Transport of Amino Acids Across the Blood-Brain Barrier. Front Physiol 11, 973 (2020).

63. Xia, R., Peng, H.-F., Zhang, X. & Zhang, H.-S. Comprehensive review of amino acid transporters as therapeutic targets. International Journal of Biological Macromolecules 260, 129646 (2024).

64. Dyer, D. L. & Said, H. M. [42] Biotin uptake in cultured cell lines. in Methods in Enzymology vol. 279 393–405 (Academic Press, 1997).

65. Rossi, A. et al. Genetic compensation induced by deleterious mutations but not gene knockdowns. Nature 524, 230–233 (2015).

66. Krill-Burger, J. M. et al. Partial gene suppression improves identification of cancer vulnerabilities when CRISPR-Cas9 knockout is pan-lethal. Genome Biology 24, 192 (2023).

67. Quevedo, R. et al. Assessment of Genetic Drift in Large Pharmacogenomic Studies. cels 11, 393–401.e2 (2020).

68. Hua, K. & Ferland, R. J. Primary cilia proteins: ciliary and extraciliary sites and functions. Cell. Mol. Life Sci. 75, 1521–1540 (2018).

69. Maes, E. et al. Determination of variation parameters as a crucial step in designing TMT-based clinical proteomics experiments. PLoS One 10, e0120115 (2015).

70. Anvarian, Z., Mykytyn, K., Mukhopadhyay, S., Pedersen, L. B. & Christensen, S. T. Cellular signalling by primary cilia in development, organ function and disease. Nature Reviews Nephrology 15, 199–219 (2019).

71. Hansen, J. N. et al. Intrinsic Diversity In Primary Cilia Revealed Through Spatial Proteomics. 2024.10.20.619273 Preprint at 10.1101/2024.10.20.619273 (2024).

72. Stefan, C. J. et al. Membrane dynamics and organelle biogenesis-lipid pipelines and vesicular carriers. BMC Biol 15, 102 (2017).

73. Scorrano, L. et al. Coming together to define membrane contact sites. Nat Commun 10, 1287 (2019).

74. Han, Y. et al. Directed Evolution of Split APEX2 Peroxidase. ACS Chemical Biology 14, 619– 635 (2019).

75. Qin, W. et al. Dynamic mapping of proteome trafficking within and between living cells by TransitID. Cell 186, 3307–3324.e30 (2023).

76. Garloff, V., Krüger, T., Brakhage, A. & Rubio, I. Control of TurboID-dependent biotinylation intensity in proximity ligation screens. Journal of Proteomics 279, 104886 (2023).

77. May, D. G., Scott, K. L., Campos, A. R. & Roux, K. J. Comparative Application of BioID and TurboID for Protein-Proximity Biotinylation. Cells 9, 1070 (2020).

78. Breslow, D. K. & Nachury, M. V. Analysis of soluble protein entry into primary cilia using semipermeabilized cells. Methods in Cell Biology (2015) doi:10.1016/bs.mcb.2014.12.006.

79. Moggridge, S., Sorensen, P. H., Morin, G. B. & Hughes, C. S. Extending the Compatibility of the SP3 Paramagnetic Bead Processing Approach for Proteomics. J Proteome Res 17, 1730– 1740 (2018).

80. Hughes, C. S. et al. Ultrasensitive proteome analysis using paramagnetic bead technology. Mol Syst Biol 10, 757 (2014).

81. Werner, T. et al. Ion coalescence of neutron encoded TMT 10-plex reporter ions. Anal Chem 86, 3594–3601 (2014).

82. Kong, A. T., Leprevost, F. V., Avtonomov, D. M., Mellacheruvu, D. & Nesvizhskii, A. I. MSFragger: ultrafast and comprehensive peptide identification in mass spectrometry-based proteomics. Nat Methods 14, 513–520 (2017).

83. Savitski, M. M., Wilhelm, M., Hahne, H., Kuster, B. & Bantscheff, M. A Scalable Approach for Protein False Discovery Rate Estimation in Large Proteomic Data Sets. Mol Cell Proteomics 14, 2394–2404 (2015).

84. Xie, Z. et al. Gene Set Knowledge Discovery with Enrichr. Current Protocols 1, e90 (2021).

85. Gambarotto, D., Hamel, V. & Guichard, P. Ultrastructure expansion microscopy (U-ExM). In Methods in Cell Biology vol. 161 57–81 (Elsevier, 2021).

86. Hansen, J. N., Rassmann, S., Stüven, B., Jurisch-Yaksi, N. & Wachten, D. CiliaQ: a simple, open-source software for automated quantification of ciliary morphology and fluorescence in 2D, 3D, and 4D images. Eur. Phys. J. E 44, 18 (2021).

87. Kowalczyk, I. et al. Neural tube closure requires the endocytic receptor Lrp2 and its functional interaction with intracellular scaffolds. Development 148, dev195008 (2021).

88. Ye, F., Nager, A. R. & Nachury, M. V. BBSome trains remove activated GPCRs from cilia by enabling passage through the transition zone. Journal of Cell Biology 217, 1847–1868 (2018).

